# Inferring the causes of noise from binary outcomes: A normative theory of learning under uncertainty

**DOI:** 10.64898/2026.03.01.708925

**Authors:** Xiaotong Fang, Payam Piray

## Abstract

Inferring the true cause of noise—distinguishing between volatility (environmental change) and stochasticity (outcome randomness)—is essential for learning in noisy environments. While most studies rely on binary outcomes, previous models are designed for continuous outcome and use ad hoc approximations to handle binary data, introducing theoretical inconsistencies and interpretational issues. Here, we develop a normative framework for inferring the causes of noise from binary feedback that remains faithful to the discrete nature of the generative process and underlying statistical structure. First, we establish a generative model using a state space approach tailored for binary outcomes and derive the corresponding hidden Markov model inference procedure. Second, we introduce a computational model combining the hidden Markov model with particle filtering to simultaneously infer volatility and stochasticity from binary outcomes. Third, we validate predictions through a 2×2 probabilistic reversal learning task with human participants, systematically manipulating both noise parameters. Results show that participants adjust their learning rates consistent with model predictions, increasing learning rates under volatile conditions and decreasing them under high stochasticity. Our theoretical and experimental results offer a principled approach for dissociating volatility and stochasticity from binary outcomes, providing insights into learning processes relevant to typical cognition and psychiatric conditions.

## Introduction

Learning under uncertainty is a fundamental challenge for humans and animals making decisions in complex environments. Whether foraging for food, interpreting social signals, or adapting to changing circumstances, organisms must constantly update their understanding of the world based on noisy observations. Critically, learners cannot directly observe the true state of the world; they only see outcomes and must infer the hidden causes that generated them. This inference is complicated by the fact that noise can arise from two distinct sources. First, the hidden state itself may change over time, a source of uncertainty known as volatility. Second, observations may be unreliable reflections of an otherwise stable hidden state, a source of uncertainty known as stochasticity. When an unexpected outcome occurs, the learner faces an inference problem: did the hidden state change, or was the outcome simply noisy? This distinction is critical because the two sources of noise call for opposite responses: if the hidden state changed, past information is outdated and learning should be fast; if the outcome was just noise, past information remains valid and learning should be slow. Correctly attributing noise to its source is therefore essential for adaptive behavior, yet this inference is challenging because both sources contribute to the same observable outcome, surprising events.

The idea that learning should adapt to the reliability of information has a long history in psychology. Theories of conditioning have emphasized that the rate of learning depends on the predictability of outcomes—from Mackintosh’s (1975) attentional theory to Pearce and Hall’s (1980) associability model. These frameworks capture the intuition that surprising or unreliable events warrant different learning responses than predictable ones. More recently, Bayesian approaches have formalized these ideas, showing that optimal learning rates depend on the noise structure of the environment (Behrens et al., 2007; Dayan et al., 2000; Nassar et al., 2010). The Kalman filter, in particular, has become influential for showing how learning rates should increase with volatility and decrease with stochasticity (Dayan & Yu, 2003; Gershman, 2015). Building on this foundation, several models have augmented the Kalman filter with inference mechanisms to estimate volatility from observations when stochasticity is assumed to be known (Behrens et al., 2007; Mathys et al., 2011). However, these models do not address the core computational problem: how learners might dissociate volatility from stochasticity when both are unknown and must be inferred from experience. We have recently developed a modeling approach that addresses this problem for continuous outcomes, allowing volatility and stochasticity to be jointly inferred from observations alone (Piray & Daw, 2021a, 2024). Normative predictions of this framework, that adaptive learners should dissociate volatility and stochasticity based on outcomes and adjust their learning rates accordingly, have been validated in our recent experimental work with human participants (Piray & Daw, 2024). However, this work focused on continuous outcomes, leaving the problem unaddressed for binary settings.

Binary outcomes are ubiquitous in both experimental paradigms and real-world settings. Probabilistic reversal learning tasks, widely used in animal and human research to study flexible learning, decision-making and their neural substrates, rely on binary feedback. Beyond the lab, many situations involve binary outcomes that may not perfectly reflect underlying states: a medical test returns positive or negative; a friendly gesture is returned or ignored; a dating request is accepted or rejected; a job application succeeds or fails. In these cases, the learner must infer a hidden state from noisy binary feedback and must determine whether unexpected outcomes reflect a change in the underlying state or noise in the observation process. Unlike continuous outcomes, which provide graded feedback, binary outcomes produce win or loss experiences that are psychologically more salient. This salience may make misattribution of noise sources particularly consequential for binary feedback, especially in psychiatric contexts. Individuals prone to self-blame, such as those with major depressive disorder, may misinterpret stochastic errors as meaningful patterns, falsely perceiving random mispredictions as evidence of failure (Abramson et al., 1978; Beck, 1970; Zahn et al., 2015). This misattribution can create a self-reinforcing cycle, where stochastic fluctuations are mistaken for systematic failures, exacerbating symptoms through increased self-blame and maladaptive belief updating. A principled framework for dissociating volatility and stochasticity from binary outcomes is therefore needed.

Existing models for binary outcomes have extended the Kalman filter framework using various approximation strategies. The most influential of these, the Hierarchical Gaussian Filter (Mathys et al., 2011), was designed to estimate volatility from binary observations when stochasticity is assumed to be known and fixed. Extending these models to the full problem, where both volatility and stochasticity must be inferred from outcomes, is not straightforward. The way these models convert continuous latent variables into binary outcome probabilities (using transformations such as sigmoid) introduces fundamental problems: stochasticity is not represented as an explicit parameter, and learning rates increase following surprising events regardless of their true source, even when volatility is zero and surprise is entirely due to stochasticity (Piray & Daw, 2020).

Here, we take a fundamentally different approach. Rather than approximating Kalman-based inference for binary data, we build on the Hidden Markov Model (HMM)—a framework specifically designed for binary latent states and binary observations. The HMM provides a tractable algorithm for binary outcomes when both volatility and stochasticity are known, analogous to the role the Kalman filter plays for continuous outcomes. Importantly, this change in perspective allows us to address the full problem: jointly inferring volatility and stochasticity from binary outcomes alone. Our contribution is three-fold: First, we establish a generative model based on the state space framework that makes explicit the latent attribution problem underlying learning and present the HMM inference procedure for this model. Second, we introduce PF-HMM, a computational model that combines HMM with particle filtering (PF) techniques (Andrieu et al., 2010; Doucet & Johansen, 2011), enabling learning of volatility and stochasticity directly from binary outcomes. Third, we present new experimental results from human participants in two 2×2 probabilistic reversal learning tasks that systematically manipulate both noise parameters.

The paper is structured to progressively build this framework. We begin by formally presenting the generative model, then review the HMM inference procedure and demonstrate through simulations how it captures learning given known volatility and stochasticity. The core computational contribution follows: integrating HMM with particle filtering to simultaneously learn both parameters from outcomes. We validate the model with two novel experimental datasets, revealing how humans differentially respond to volatility and stochasticity when learning from binary outcomes.

## Results

### State space models

We begin by considering a relatively general class of probabilistic models known as state space models in control theory (Ljung, 1998) and Markov models in machine learning (Bishop, 2006). In these models, a hidden “state” variable, *x*_*t*_, evolves over discrete time steps, and outcomes, *o*_*t*_, are generated based on this hidden variable but corrupted by noise. Specifically, the model’s structure defines two key conditional dependencies: *p*(*x*_*t*_|*x*_*t*−1_, *v*), where *v* is the volatility parameter governing the noise in the state transition process, and *p*(*o*_*t*_|*x*_*t*_, *s*), where s is the stochasticity parameter controlling noise in the outcome process. For inference in this model, the observer must compute the posterior probability *p*(*x*_*t*_|*o*_1_, *o*_2_, …, *o*_*t*_) given all outcomes up to the current point. Importantly, this state space modeling approach can be implemented with either continuous or binary variables. When the hidden and observed variables are continuous, inference leads to the Kalman filter, while with binary variables, it leads to the HMM. We provide a formal treatment of this framework in Appendix A; here, we focus on the key intuitions.

### Kalman-based hierarchical models

When both the hidden state and outcomes are continuous, and the conditional dependencies are Gaussian, optimal inference is tractable and given by the Kalman filter. The Kalman filter provides a closed-form solution in which the posterior mean (the best estimate of the hidden state) is updated by a delta-rule with a dynamic learning rate (Appendix A.2). Crucially, this learning rate depends on both volatility and stochasticity: it increases with volatility and decreases with stochasticity, reflecting how quickly an organism should revise its expectations in the face of prediction errors. This framework has become foundational in statistical theories of conditioning, with the Kalman gain corresponding directly to the associability term in classical conditioning models (Dayan et al., 2000; Dayan & Long, 1998; Dayan & Yu, 2003). It has helped explain numerous phenomena in the conditioning literature, including blocking, trial-order effects, auto-shaping, and latent learning (Dayan & Yu, 2003; Gershman, 2015; Kakade & Dayan, 2002). More broadly, this Bayesian approach formalizes the idea that animals infer causal relationships between variables, including unobservable ones, and estimate uncertainty surrounding these relationships, allowing them to weigh new evidence against prior knowledge based on their relative reliability (Courville et al., 2006; Gershman et al., 2010; Piray & Daw, 2021a).

Here, we will focus on hierarchical modeling approaches built on the Kalman filter framework, which extends learning to account for volatility, particularly in the context of binary outcomes. These models typically augment the standard generative model by treating volatility as a dynamic variable that evolves according to its own conditional transition probability, *p*(*v*_*t*_|*v*_*t*−1_). This extension, while theoretically elegant, introduces a significant computational challenge: the inference problem is no longer analytically tractable, necessitating various forms of approximate inference. Several notable approaches have addressed this challenge. In their seminal work, Behrens and colleagues employed a brute force numerical integration method to perform approximate inference in this hierarchical framework (Behrens et al., 2007, 2008). A more computationally sophisticated solution was subsequently introduced by Mathys and colleagues, who developed a parametric variational approach (Iglesias et al., 2013; Mathys et al., 2011). In their model, the posterior distributions over both *x*_*t*_ and *v*_*t*_ are approximated by Gaussian distributions through an explicit hierarchical update procedure. This procedure first computes the mean and variance of the approximate posterior for *x*_*t*_ and then calculates the mean and variance of the approximate posterior for *v*_*t*_. Specifically, Mathys et al. assumed that the transition probability for *v*_*t*_ follows a Gaussian distribution and implemented an exponential transformation function to appropriately scale the variance of the transition probability before incorporating it into the next level of the hierarchy (Iglesias et al., 2013; Mathys et al., 2011). We have previously proposed a slightly different formulation for the transition probability of volatility, utilizing Gamma distribution instead (Piray & Daw, 2020).

Particularly relevant to our present discussion is how these models have been adapted for binary outcomes. These models modified the conditional dependency of the continuous generative model to accommodate binary outcomes, using *p*(*o*_*t*_|*x*_*t*_) = Bernoulli(σ(*x*_*t*_)), where σ(*x*) = 1/(1 + exp(−*x*)) is the sigmoid function. This formulation has an important conceptual implication: it effectively eliminates an explicit stochasticity parameter. This does not mean the absence of stochasticity in the model; rather, stochasticity exists but becomes conflated with other parameters. For example, even with zero volatility and an initial hidden variable value of 1, the probability of observing a positive outcome (1) is 0.73. Thus, outcomes remain stochastically generated despite the lack of an explicit stochasticity parameter. To implement inference with binary outcomes, these models have employed various well-established approximation strategies, such as Taylor expansions (Mathys et al., 2011) and moment matching (Piray & Daw, 2020) techniques. Despite their technical differences, these approaches share a common foundation: they preserve the underlying generative structure of the Kalman filter, allowing them to derive approximate inference methods with update equations that closely resemble the standard Kalman update rules. However, as we have formally shown previously (Piray & Daw, 2020), this approach introduces a significant theoretical problem. The learning rate—defined as the ratio of prediction update to prediction error—exhibits a problematic relationship with volatility: it increases dramatically following large, surprising events (i.e., large unsigned prediction errors), regardless of the true volatility level, and even when volatility is zero (Supplementary Figure 1). This spurious effect stems directly from combining the Kalman filter with the sigmoid transformation used to convert the continuous hidden variable *x*_*t*_ into a probability (see Appendix B for a formal treatment). This limitation is particularly problematic for experimental designs that manipulate volatility, such as reversal learning tasks with distinct low and high volatility blocks. In high-volatility blocks, where contingencies change frequently, large surprises naturally occur more often, creating an artificial correlation between volatility and the implied learning rate. But the problem runs deeper: even when volatility is fixed, the learning rate increases with unexpected outcomes that are entirely due to stochasticity. This fundamental confound underscores the need for computational models that properly disentangle these distinct sources of uncertainty.

### Hidden Markov Model

In contrast to previous approaches, we develop a generative model that more faithfully represents the inherent structure of binary environments. Rather than adapting continuous-variable frameworks to binary data, we directly exploit the established architecture of state space models specifically designed for binary hidden causes—an approach with strong foundations in machine learning (Bishop, 2006; Wainwright & Jordan, 2008), for which the inference is tractable and given by the HMM (Ghahramani, 2001; Rabiner, 1989). The HMM has broad applications across the biological sciences, including areas such as genetics (Burge & Karlin, 1998; Krogh et al., 2001), and ecology (McClintock et al., 2020). HMMs have also been widely used in neuroscience for analysis of neural data (Baldassano et al., 2017; Bolkan et al., 2022; Calhoun et al., 2019; Vidaurre et al., 2018; Wiltschko et al., 2015). However, unlike the Kalman filter, HMMs have rarely been used as a computational model to explain phenomena related to conditioning and learning, though they have occasionally been used to fit human experimental data (Hampton et al., 2006; Schlagenhauf et al., 2014).

Here, we present a generative model consisting of a binary hidden state and binary outcomes, with two distinct sources of noise affecting the process at different levels. We provide the main equations below; see Appendix A.3 for the full derivation. Specifically, the model centers on a binary hidden state variable, *x*_*t*_, which can take only two values (0 or 1) and evolves over discrete time steps according to probabilistic rules. From this hidden state, binary outcome, *o*_*t*_, is generated but potentially corrupted by outcome noise (Figure 1). This framework allows us to explicitly model both the volatility of the environment and the stochasticity of outcomes as independent sources of noise with separate parameters.

In particular, the hidden variable evolves according to the following rule:

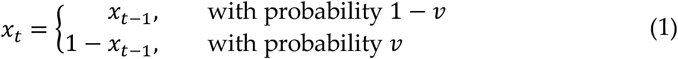

where 0 ≤ *v* ≤ 0.5 is the volatility parameter governing the degree of noise in this process. If *v* = 0 there is no diffusion noise, and the hidden state never changes. If *v* = 0.5, there is an equal probability that the hidden state switches or stays the same on every trial, indicating that the noise is maximum. This means that the conditional transition probability is given by a Bernoulli distribution:

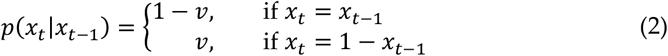

The binary outcome is noisily generated based on the hidden state, with the amount of noise governed by the stochasticity parameter:

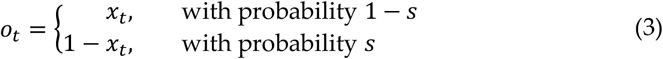

where 0 ≤ *s* ≤ 0.5 is the stochasticity parameter governing the degree of noise. The outcome switches with respect to the hidden variable with a probability given by *s*. If *s* = 0, there is no outcome noise, and the outcome always follows the hidden state. If *s* = 0.5, on the other hand, there is equal probability that the outcome switches or follows the hidden state, essentially generating outputs completely randomly regardless of the hidden state. Thus, the conditional outcome probability is also given by a Bernoulli distribution:

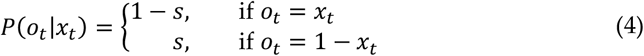

To illustrate, consider a typical probabilistic reversal learning task, where participants learn that one option is rewarded 80% of the time while the other is rewarded 20%, and these contingencies periodically reverse (Cools et al., 2002; Costa et al., 2015; Izquierdo et al., 2017). In our framework, the reversal frequency corresponds to volatility (*v*), while the 80/20 reward schedule corresponds to stochasticity (*s*). Critically, from the participant’s perspective, both sources of noise produce the same observable result: an unexpected outcome. When a participant chooses the supposedly better option but receives no reward, this could mean the contingencies have reversed (volatility), or it could simply be the 20% chance of no reward from a still-correct option (stochasticity). Disentangling these possibilities is the core computational challenge our framework addresses.

For our binary state space model, the inference process—computing the posterior probability distribution *p*(*x*_*t*_|*o*_1_, *o*_2_, …, *o*_*t*_)—is analytically tractable with simple update equations within the HMM framework (Ghahramani, 2001). The HMM framework gives exact, closed-form equations for how beliefs should be optimally updated in light of new outcomes.

The posterior distribution maintains a Bernoulli form at each time step: *p*(*x*_*t*_|*o*_1_, *o*_2_, …, *o*_*t*_) = Bern(*x*_*t*_|*r*_*t*_) where *r*_*t*_ represents the probability that the hidden state equals 1 given all outcomes up to and including time *t*. This parameter *r*_*t*_ is updated recursively through a set of equations that incorporate the previous belief state, volatility parameter, stochasticity parameter, and current outcome. Specifically, the update process involves three steps:

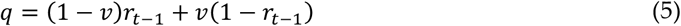

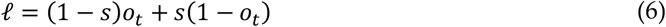

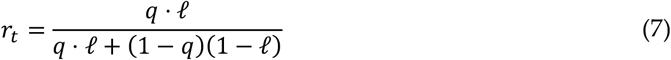

with *r*_0_ = 0.5. These equations have clear interpretations:

- The intermediate variable *q* equals *p*(*x*_*t*_ = 1|*o*_1_, *o*_2_, …, *o*_*t*−1_), representing our belief about the current state before seeing the current outcome. This quantity incorporates both the previous belief (*r*_*t*−1_) and the probability of state transitions (governed by volatility *v*).
- The variable *l* equals *p*(*o*_*t*_|*x*_*t*_ = 1), representing the likelihood of observing the current outcome *o*_*t*_ if the hidden state is 1. This likelihood depends on the stochasticity parameter *s*, which determines how reliably the outcome reflect the state.
- The final equation for *r*_*t*_ is an application of Bayes’ rule, combining the prior belief *q* with the likelihood of the outcome, *l*, to compute the posterior probability.

This recursive Bayesian update scheme provides a computationally efficient method for tracking belief states as new outcomes arrive. The equations maintain mathematical elegance while capturing how both volatility and stochasticity affect the learning process in binary environments (for full derivations see Appendix A.3).

Unlike the Kalman filter, there is no explicit learning rate variable in the update rules of the HMM. Nevertheless, the learning rate, defined as the ratio of prediction update to prediction error serves as an informative descriptive measure of learning, even if the underlying process does not involve an explicit error-driven update. Specifically, the HMM provides a prediction on each trial (i.e., *r*_*t*_ in Eqs. 5-7), allowing for the computation of both a prediction error and an update. As a result, the learning rate can be quantified as (*r*_*t*_ − *r*_*t*−1_)/(*o*_*t*_ − *r*_*t*−1_). This approach is similar in practice to how prior studies using binary outcomes have typically assessed learning.

We thus investigated how volatility and stochasticity influence the learning rate in the HMM. To do this, we defined a 2 × 2 prototypical design, which includes a systematic manipulation of both volatility and stochasticity (Figure 2A). This results in four blocks of binary time series, where values of volatility and stochasticity are either small or large. Next, we simulated the HMM for each block separately using the true values of volatility and stochasticity. This allowed us to quantify the learning rate by regressing the trial-by-trial update in predictions against prediction error across blocks. Block-specific intercepts were also included in the regression analysis. This method yields the learning rate per block. Notably, this approach reveals the theoretically optimal learning rate (i.e., ideal observer learning rate), since the HMM is the optimal inference model for this generative process and our simulations use the true parameter values for volatility and stochasticity. This simulation analysis (Figure 2B) revealed that the learning rate for binary outcomes follows the intuitive predictions: higher volatility increases the optimal learning rate, indicating rapid adaptations to new information. Conversely, higher stochasticity decreases the optimal learning rate, indicating the need for having a larger window for averaging outcomes, as each individual outcome is less reliable.

**Figure 1.**
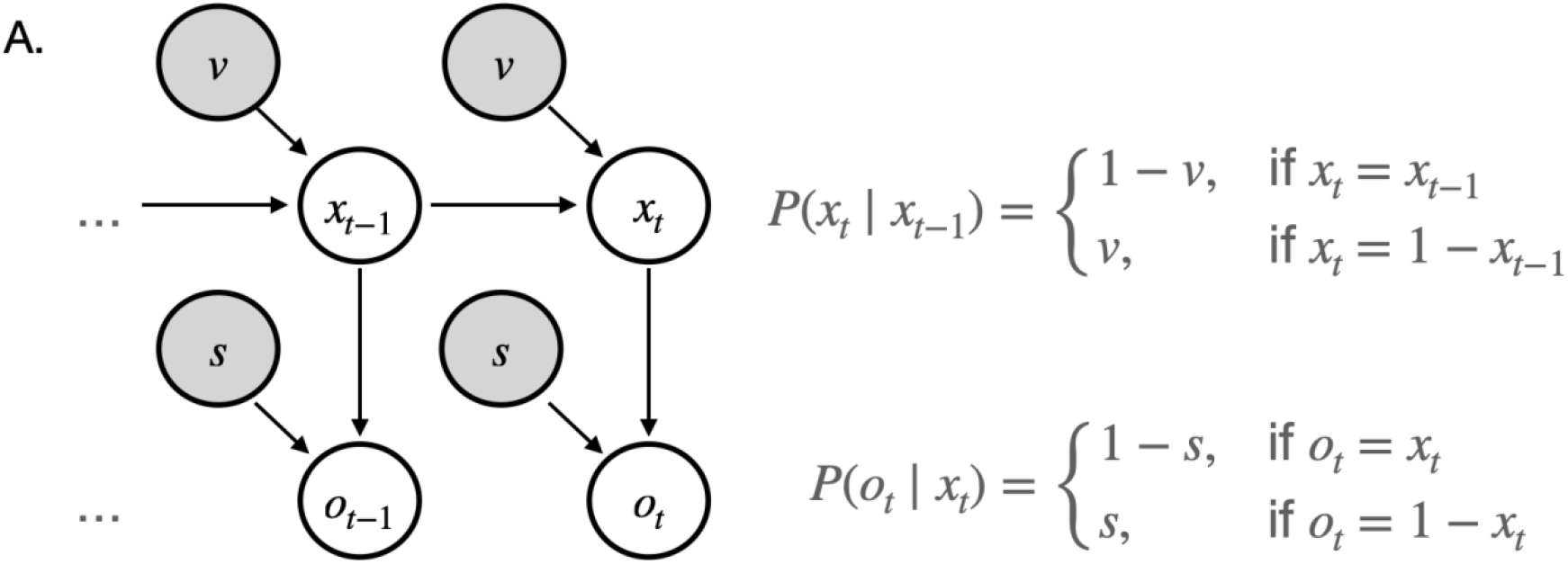
Graphical model of Hidden Markov Model. The model infers hidden state (*x*_*t*_) by updating predictions based on observed outcomes (*o*_*t*_), where volatility (*v*) controls the probability of state transitions and stochasticity (*s*) determines the noise in outcome generation. In the model, *v* and *s* are treated as constants, determining the volatility of the hidden state and the level of stochasticity in outcomes relative to it, respectively.

**Figure 2.**
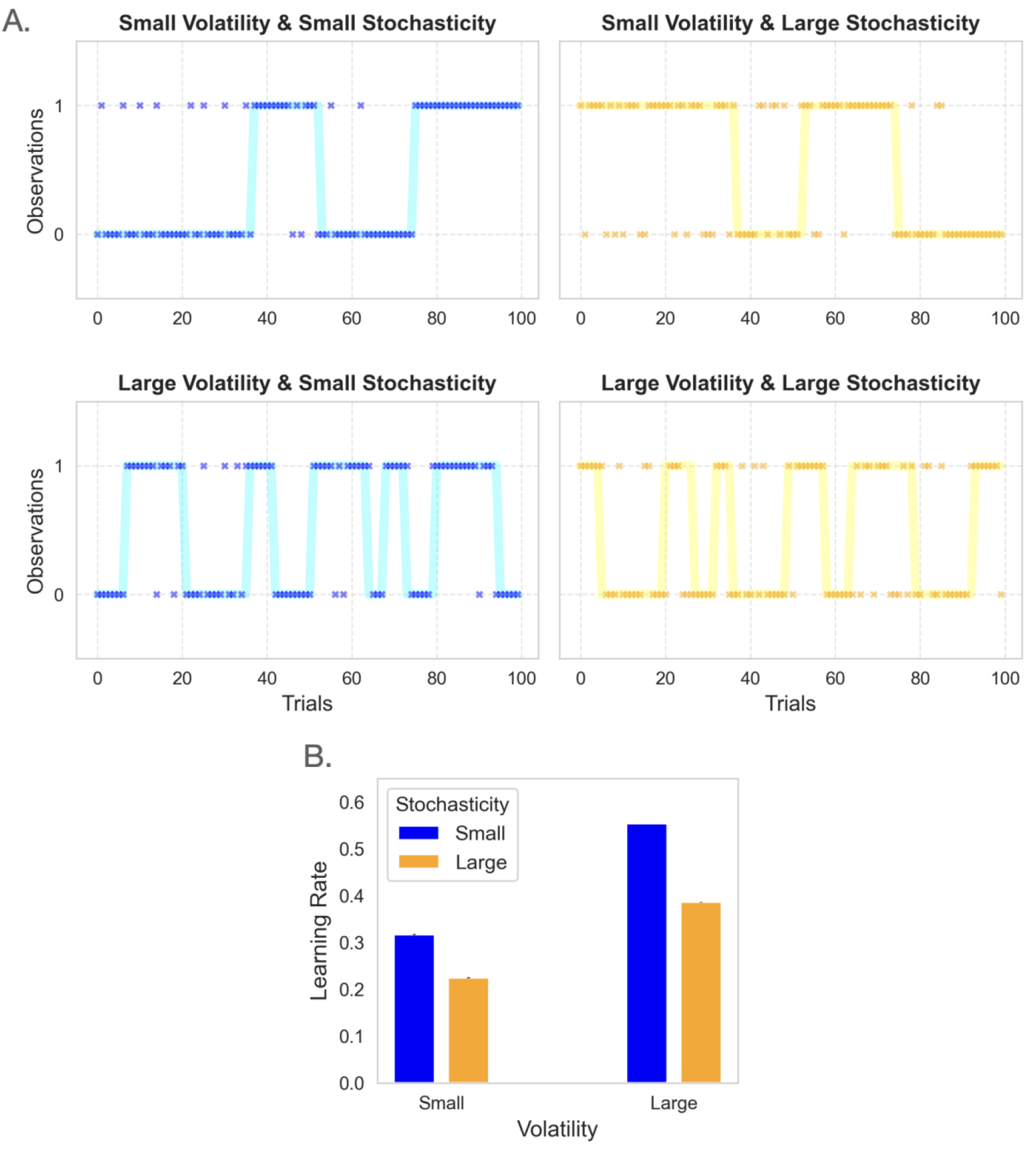
HMM simulation analysis. (A) Examples of generated time series with volatility levels of 0.025 and 0.1, and stochasticity levels of 0.125 and 0.25. The solid lines represent the hidden state, whose variability depends on volatility levels. Individual data points are marked separately, with their variations relative to the hidden state determined by the stochasticity level. (B) Learning rates quantified by the optimal HMM—given the true values of volatility and stochasticity—vary systematically with both factors. The simulation was repeated 1000 times and the means and standard errors are plotted. The model suggests that volatility and stochasticity exert opposing effects: higher volatility increases the learning rate, while higher stochasticity decreases it.

### PF-HMM: a model for inferring the cause of noise based on binary outcomes

The HMM model presented above provides an optimal algorithm for recursively computing the posterior over the hidden state when new outcomes arrive. However, this algorithm, similar to the Kalman filter, relies on the critical assumption that the values of volatility and stochasticity are known and given. In real-world environments, neither of these values is known a priori, and both must be estimated from observations. This presents a fundamental challenge, especially when both volatility and stochasticity are unknown and potentially fluctuating over time.

To address this challenge, we extend the state space model presented above to include both volatility and stochasticity as dynamically changing random variables. Specifically, we expand the state space model of Figure 1 by assuming that *v*_*t*_ and *s*_*t*_ evolve according to independent diffusion processes (Figure 3B). Each variable drifts smoothly over time around its previous value, controlled by separate diffusion parameters. Importantly, this formulation allows the variables to fluctuate over time without being biased to drift in any direction. The formal mathematical definitions of these diffusion processes have been detailed in Appendix A.4.

**Figure 3.**
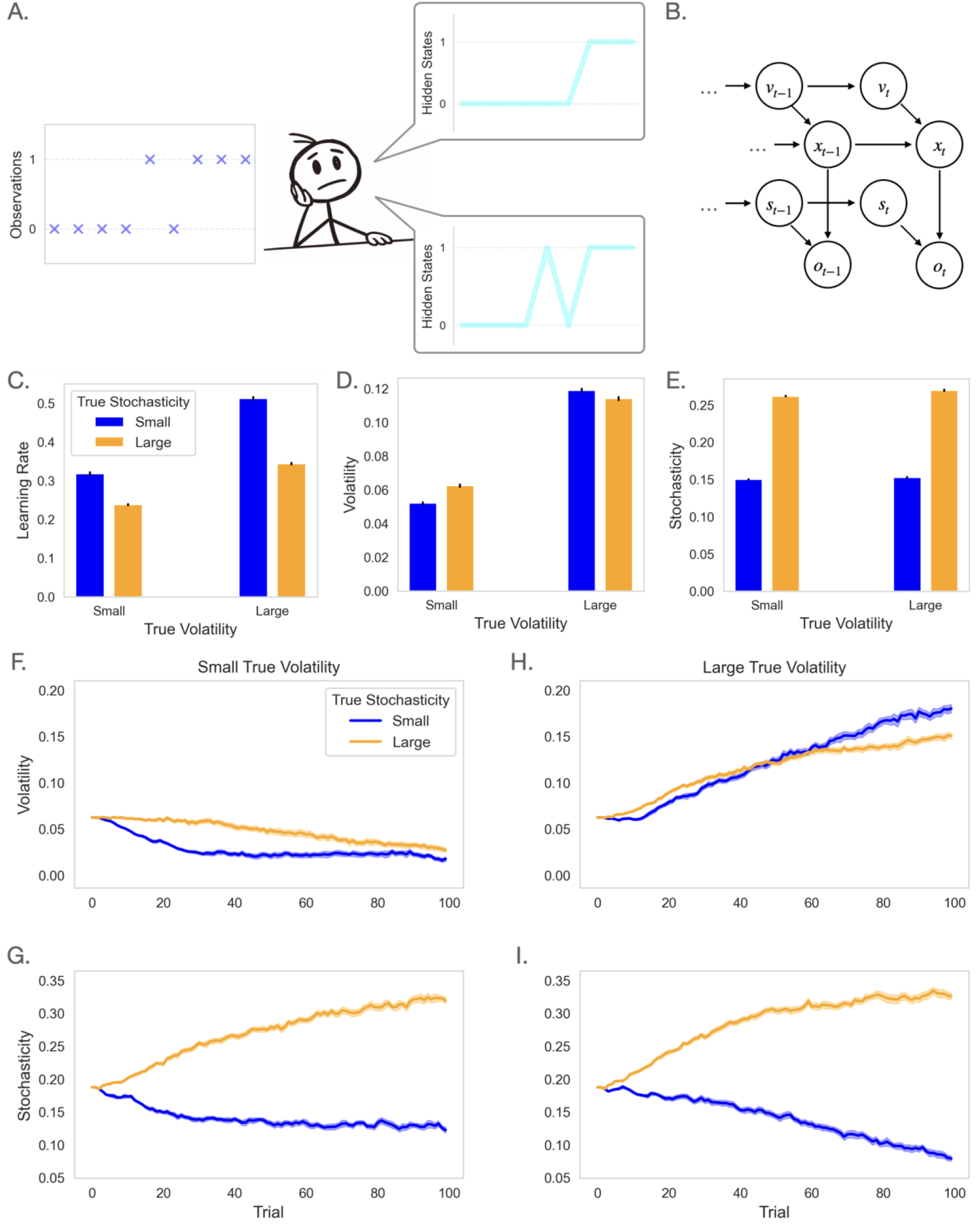
PF-HMM simulation analysis. (A) In the real world, one must infer the hidden variable based solely on observations, which leads to trial-by-trial attribution (or misattribution) of experienced noise to either volatility or stochasticity. The same observed outcome sequence can arise either from frequent hidden-state changes (high volatility, low stochasticity) or from a stable state corrupted by outcome noise (high stochasticity, low volatility). Because the latent state is unobserved, learners must infer the source of variability, creating a risk of misattribution that leads to different learning-rate adjustments. (B) The structure of the (generative) model: The binary outcome on trial *t*, denoted *o*_*t*_, is generated from a binary hidden state *x*_*t*_, corrupted by independent noise whose magnitude is determined by the level of stochasticity, *s*_*t*_. The hidden state itself evolves from its previous value, *x*_*t*−1_, corrupted by another source of noise governed by the volatility parameter, *v*_*t*_. Both *v*_*t*_ and *s*_*t*_ also evolve over time, changing noisily based on their values in the previous trial. The learner’s task is to infer the current hidden state, as well as volatility and stochasticity, based on the sequence of observed outcomes. (C-I) PF-HMM was simulated using the same 2 × 2 design as in Figure 2B. The model’s learning rate is influenced by both true volatility and stochasticity, which have opposite effects: higher volatility increases the learning rate, while higher stochasticity decreases it (C). Estimated volatility captures differences in true volatility (D), and estimated stochasticity captures differences in true stochasticity (E). Panels F-G show volatility and stochasticity estimates from the model for blocks with low true volatility, while panels H–I show estimates for blocks with high true volatility. The simulation was conducted on the same 1000 time series used in the analysis shown in Figure 2. Error bars in panels C–E represent the standard error of the mean, and those in panels F–I represent the standard error of the median.

Since this model no longer permits exact inference, we turn to approximate inference techniques. We employ a PF approach, a standard and efficient method broadly applicable to such problems (Andrieu et al., 2010; Doucet et al., 2001). The key idea is to maintain a population of candidate hypotheses (particles), each representing a possible combination of volatility and stochasticity values. As outcomes arrive, particles that better explain the data receive more weight, allowing the model to track the noise parameters over time.

Importantly, the model leverages a key insight: when provided with samples of volatility and stochasticity parameters, we can conduct tractable inference for the hidden state through the HMM framework. This combined strategy is known as Rao-Blackwellized PF (Doucet et al., 2000; Doucet & Johansen, 2011), which efficiently handles the computational challenges of our the extended generative model. In sum, the new PF-HMM model estimates *v*_*t*_ and *s*_*t*_ using PF, while inference for *x*_*t*_ is conducted via HMM, conditional on these estimated parameter values.

The algorithm operates in three distinct steps on each trial. The first step involves prediction, where individual particles transition forward in time according to the generative dynamics for volatility and stochasticity. This gives rise to a set of sampled values for these variables. The second step evaluates how well each particle’s predictions match the observed outcome.

Particles that better predict the outcome receive higher weights, while those that predict poorly are down weighted. As is customary in particle filtering methods, systematic resampling is applied during this phase if the effective number of particles drops below a certain threshold (Douc & Cappe, 2005). The third step applies the HMM to integrate the new outcome with previous estimates, updating the belief about the hidden state for each particle. The expected value for all random variables at each trial is computed as the weighted average across all particles. The complete formal algorithm is given in Appendix A.4.

Next, we tested this model using the same 2 × 2 paradigm described above. In this paradigm, the true values of volatility and stochasticity are fixed within each block and only change between blocks. However, the model has no prior knowledge of this structure. Instead, the initial values for volatility and stochasticity are set to the midpoint between the low and high true values used in the experiment, and the diffusion parameters for *v*_*t*_ and *s*_*t*_ are both set to 0.02. Thus, this approach allows us to examine how well the model can adapt to the different volatility and stochasticity conditions across blocks without being explicitly informed about the block structure or the true parameter values. By keeping the model’s parameterization constant across all conditions, we can observe its ability to infer the underlying environmental statistics purely from outcomes.

Figure 3 shows the results of this analysis. We first quantified the learning rate per block. As shown in Figure 3C, the model’s learning rate exhibits a pattern very similar to the optimal learning rate presented in Figure 2B: learning rates increase with true volatility and decrease with true stochasticity. Notably, this occurs despite the fact that the model has no prior knowledge of the true volatility and stochasticity values, requiring it to estimate these parameters solely from outcomes.

Furthermore, examining the model’s estimates of volatility and stochasticity reveals it performs remarkably well in reconstructing these underlying parameters. Importantly, the model’s estimate of volatility shows minimal sensitivity to manipulations of true stochasticity, while its estimate of stochasticity remains largely unchanged across different levels of true volatility. This selective sensitivity demonstrates that the model can successfully attribute experienced noise to its actual source, distinguishing between environmental change (volatility) and observational noise (stochasticity).

### Ablated variants of PF-HMM

Our final set of models comprises ablated models of volatility and stochasticity. Previous simulation analyses revealed a crucial aspect of the model: volatility and stochasticity inferences are inherently interrelated. Although these are independent sources of noise from the perspective of the generative model, they become dependent for the purpose of inference (i.e., in the eyes of the observer). This is exactly why dissociating volatility and stochasticity is computationally challenging; both factors increase experienced noise (i.e., variability in outcomes; Figure 4). Separating these contributions requires a process of credit assignment, or as it is called in the Bayesian literature, “explaining away,” where competing explanations for outcomes are weighed against each other. A pathological model that ignores one factor incorrectly attributes noise from that factor to the other. This dynamic may be particularly relevant to neurological and psychiatric conditions that selectively impair the processing of volatility or stochasticity. Our model predicts that such impairments would not only reduce learning rate modulation but could also lead to paradoxical reversals, where volatility and stochasticity are mistakenly substituted for one another due to a breakdown in the explaining away mechanism.

**Figure 4.**
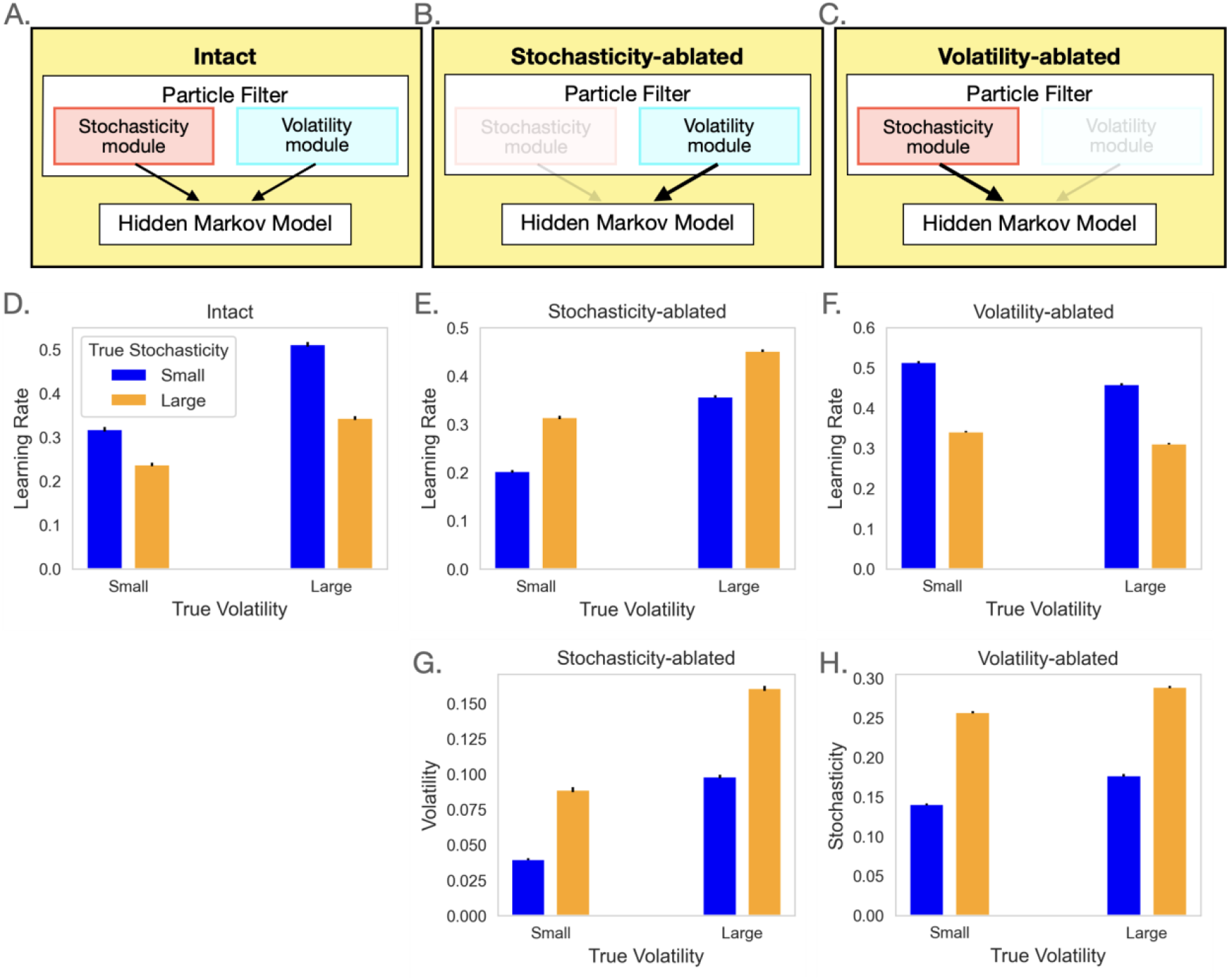
Ablated variants of PF-HMM simulation analysis. (A) In the intact model, stochasticity and volatility compete to explain the observed noise arising from the mismatch between the HMM component’s predictions and the actual outcomes. The intact model produces behaves optimally by inferring the source of noise based on observed outcomes. (B-C) Two distinct ablated models exhibit different pathological behaviors: ablating the stochasticity module results in a fixed level of stochasticity across blocks, causing noise due to stochasticity to be misattributed to volatility. In contrast, ablating the volatility module results in a fixed level of volatility across blocks, which leads to volatility-related noise being misinterpreted as stochasticity. (D-H) The intact model’s learning rate is the same as in Figure 3C and is reproduced here for comparison (D). For both ablated models, the effect of ablating does not merely eliminate the corresponding influence on the learning rate but instead causes a reversal. Specifically, the stochasticity-ablated model exhibits an elevated learning rate in response to increasing stochasticity (E). On the contrary, the volatility-ablated model shows a reduced learning rate as volatility rises (F). This reversal occurs because the noise from the ablated factor is misattributed to the remaining module. (G) The stochasticity-ablated model incorrectly infers increased volatility in highly stochastic environments. (H) Conversely, the volatility-ablated model incorrectly attributes volatility to stochasticity, resulting in an inflated stochasticity estimate in more volatile environments. The simulation was conducted on the same 1,000 time series used in the analysis shown in Figures 2–3. Error bars reflect the standard error of the mean computed over 1,000 simulations.

Figure 4 illustrates this effect using our 2 × 2 design, highlighting two distinct ablated models. In the stochasticity-ablated model, all experienced noise is attributed to volatility, causing a persistent increase in learning rate regardless of the actual source of noise (Figure 4E). Conversely, in the volatility-ablated model, all noise is attributed to stochasticity, resulting in a decrease in learning rate irrespective of the true source of noise (Figure 4F). In other words, a model lacking one noise parameter inevitably misattributes experienced noise to the remaining one, leading to systematic errors in learning. This maladaptive pattern of misattributions is also evident in the volatility and stochasticity estimates of both ablated models: In the stochasticity-ablated model, volatility estimates always increase regardless of the true source of noise (Figure 4G). Similarly, in the volatility-ablated model, stochasticity estimates always increase, irrespective of the actual source of noise (Figure 4H).

These ablated variants of the model offer a promising framework for understanding disrupted learning in clinical populations. This is particularly important in the context of psychiatric disorders, such as anxiety disorders, depression, impulse control disorders, and schizophrenia. Although maladaptive processing of uncertainty has been associated with all these disorders, it remains unknown how these same computational mechanisms might influence many mental illnesses transdiagnostically. Our framework can provide valuable insights into these shared computational dysfunctions across different psychiatric conditions.

### Human participants infer the source of noise from binary outcomes

We conducted a behavioral experiment to examine whether humans dissociate between volatility and stochasticity from binary outcomes. Following our prototypical paradigm, we designed a task to study learning from binary outcomes in a 2 × 2 setting where both factors were systematically manipulated. The task consisted of four blocks, each with distinct fixed parameters for true volatility and stochasticity, though participants were not informed of these values or their order across blocks. We recruited 73 participants (34 female, age range: 18-66, median age: 35) through the online platform Prolific.

We developed an engaging cover story (Figure 5A) in which participants predicted which side of a beach a sea lion would visit, based on where treasures had been discovered on previous trials. Crucially, participants could not observe the sea lion directly; they could only use the treasure locations to guide their predictions. They were told that the sea lion tended to stay on the same side of the beach but could suddenly switch (volatility). They also learned that ocean waves could independently carry treasures to the opposite side (stochasticity). The instructions carefully emphasized that volatility and stochasticity were independent sources of uncertainty, stemming from the sea lion’s behavior patterns and wave effects, respectively. Participants were also informed they would interact with four different sea lions on four different beaches. Each sea lion–beach pair constituted one block of 40 trials, although participants were not informed about the length of the blocks. This experimental design enabled us to measure participants’ behavioral responses to systematic manipulations of both true volatility and true stochasticity (See Supplementary Figure 2 for the timeseries used in the task).

**Figure 5.**
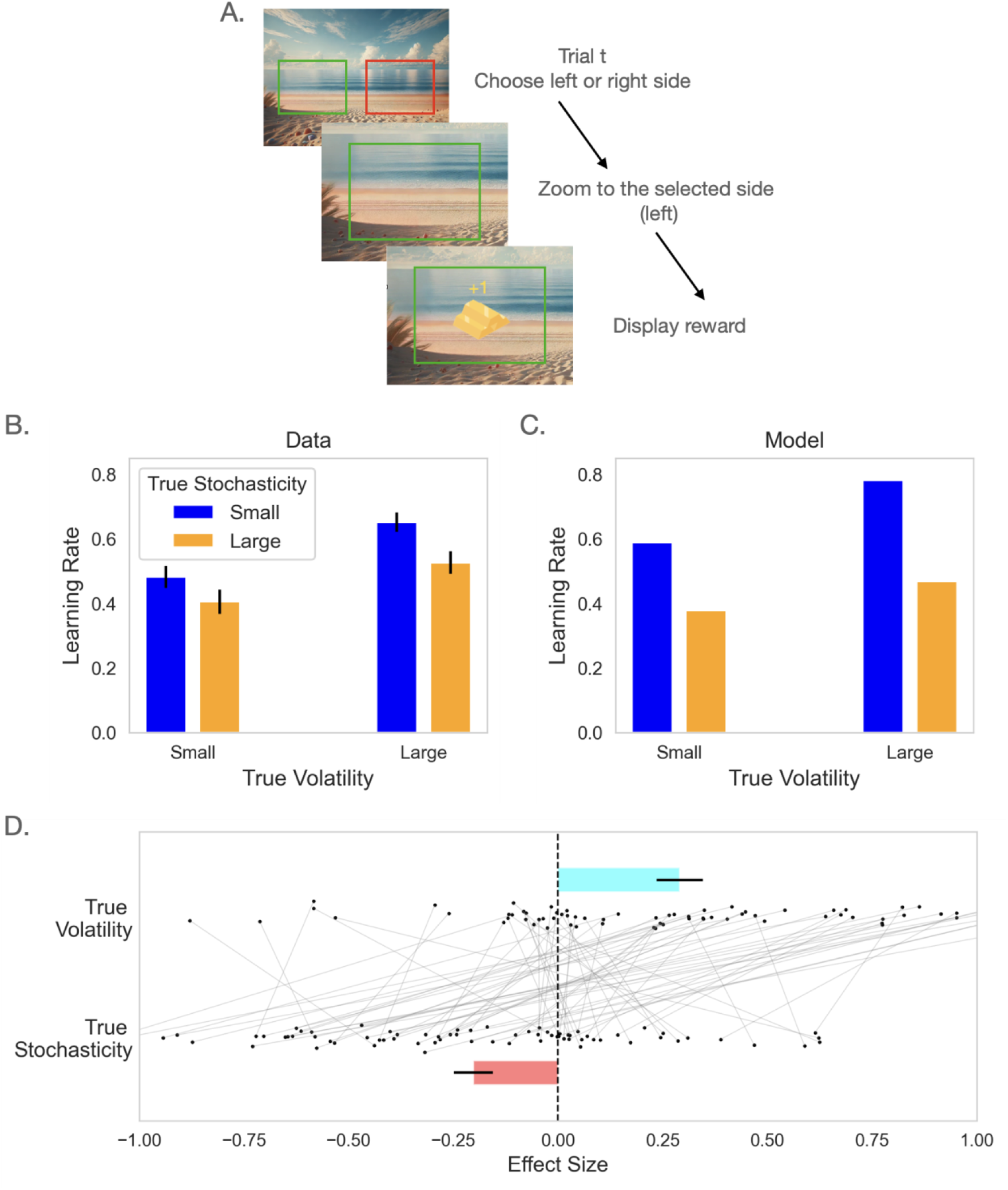
Experimental findings. (A) On every trial, participants choose one side of the beach to find treasures brought by a sealion not visible to them. Participants can use left or right arrow key to choose one side when they see the whole view of the beach. Then the screen will zoom in to that side and display either reward or no reward. The task has four blocks with a 2 × 2 factorial design, manipulating both true volatility and true stochasticity. In the illustration, the left side with green box is chosen (the boxes are not displayed in the actual task). (B) Model-neutral analysis of behavioral data. Mean learning rate per block is plotted (n = 73), which calculated by fitting the blockwise HMM model. The results show strong main effects of both factors: the learning rate increases with higher true volatility and decreases with higher true stochasticity. Note that this analysis is theory-neutral with respect to changes in learning rate across blocks, as the HMM does not impose any assumptions about how volatility and stochasticity parameters vary across blocks. (C) Learning rate from fitting the PF-HMM model on subject data shows similar effects to model neutral analysis. (D) Main effects of both factors on the learning rate coefficient have been plotted for all participants. The effect size of true volatility is calculated by learning rate of two high volatility blocks minus learning rate of two low volatility blocks. Effect size of true stochasticity is calculated similarly. Mean and standard error of the mean are plotted in B-D.

We first used a blockwise HMM to analyze choice data and quantify the learning rate per block. The model was fitted to participants’ responses, quantifying two parameters per block: a volatility parameter, *v*, and a stochasticity parameter, *s*. We used maximum a posteriori estimation with wide priors to fit these three parameters to choice data (Piray, Dezfouli, et al., 2019); see Methods for further details. Notably, this analysis is theory-neutral with respect to inference about volatility and stochasticity, as the HMM does not impose any assumptions about how these factors should vary across blocks. Therefore, any observed differences in fitted parameters or learning rates across blocks should be attributed solely to differences in participants’ behavioral responses rather than any model-driven assumptions. We also conducted a recovery analysis in which the blockwise HMM model was simulated using different sets of known parameters. The resulting time-series data were then analyzed using the same model-fitting procedure. This analysis demonstrated that the model’s parameters are almost perfectly recoverable using this approach (Supplementary Table 1 and Supplementary Figure 3).

Using the best-fitted parameters (Supplementary Table 2), we generated predictions and quantified the learning rate per block (Figure 5B). We then analyzed these learning rates across all participants. Our analysis revealed a significant positive main effect of true volatility on learning rate (*t*(72) = +5.286, *P* < 0.001), indicating that participants appropriately increased their learning rates in response to higher volatility. Conversely, we found a significant negative effect of true stochasticity on learning rate (*t*(72) = −4.302, *P* < 0.001), demonstrating that participants decreased their learning rates when true stochasticity increased, consistent with normative predictions from the model. No significant interaction between the two factors was found (*t*(72) = −1.242, *P* = 0.22). The regression analysis also revealed block-level effects that were independent of prediction error; see Supplementary Table 3. These results extend previous work demonstrating effects of volatility on learning from binary outcomes by additionally revealing a negative effect of stochasticity on learning.

Importantly, fitting PF-HMM to participants response exhibits a very similar pattern of effects (Figure 5C). A practical challenge in fitting PF models (and related Monte Carlo sampling approaches) is that the likelihood function is not differentiable with respect to the generative parameters, which has limited the application of these models in psychology. To address this, we followed the procedure that we have used in tasks with continuous observations (Piray & Daw, 2024), and adopted an optimization scheme based on Gaussian processes that is well suited for non-differentiable likelihoods (Gelbart et al., 2014; Snoek et al., 2012). We validated this approach through a parameter recovery analysis: the model was simulated with known parameter values, and the resulting data were subjected to the same fitting procedure. This analysis confirmed that the model’s parameters are reliably recoverable (Supplementary Table 4, Supplementary Figure 4). This suggests that participants dissociate volatility and stochasticity based solely on observed binary outcomes.

### Human responses are slower when identifying the true source of noise is more difficult

The previous analysis demonstrates that human participants can dissociate volatility and stochasticity by observing outcomes alone, in line with predictions from the PF-HMM. Here, we aim to further elucidate the cognitive and computational mechanisms underlying this dissociation by examining whether the PF-HMM also accounts for a related but statistically independent aspect of behavior—namely, response times.

Our modeling approach offers insight by predicting that identifying the source of noise becomes more difficult when particles are indistinguishable. This indistinguishability arises when most or all particles assign similar likelihoods to a given outcome, resulting in low variability across per-particle likelihoods. Under these conditions, the model struggles to determine which particles best explain the observation, as all provide similarly good fits. This ambiguity makes it harder to accurately attribute the observed noise to its true source— volatility or stochasticity. In contrast, high variability across particles suggests that some particles account for the outcome much better than others, enabling more confident inference about the underlying cause of the noise.

To test these predictions, we analyzed response times, anticipating that they might reveal additional signatures of the underlying computational process. Specifically, we hypothesized that responses would be slower when the core computational task—identifying the source of noise—is more difficult, which occurs when particles are indistinguishable (i.e., when particle likelihood variability is low). We quantified trial-by-trial particle likelihood variability as the standard deviation of per-particle likelihoods across particles and performed a regression analysis examining the relationship between response times and particle likelihood variability. To control for potential confounding effects, we included additional regressors: average particle likelihood, estimated volatility, estimated stochasticity, and an accuracy regressor indicating whether the response matched the outcome, ensuring that any observed effects are not due to a response time and accuracy trade-off. Model predictions were generated using the median fitted parameters across subjects (For more details see Methods).

This analysis revealed a highly significant negative relationship between response time and particle variability (*t*(72) = −3.533, *P* = 0.001; Figure 6). These results suggest that when variability across particles is low, participants respond more slowly, likely reflecting the increased difficulty of accurately identifying the source of noise due to greater indistinguishability in how particles fit the observed outcome (See also Supplementary Table 5).

**Figure 6.**
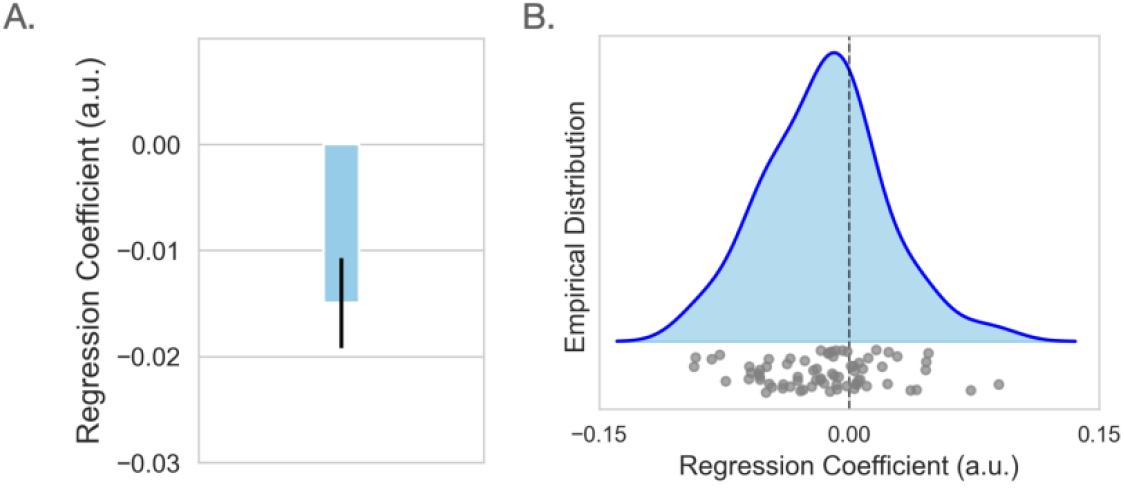
Computational mechanisms underlying how participants infer the source of noise. (A) The regression coefficient from an analysis of participants’ response times (in seconds) against particle likelihood variability (*PLV*) fitted to each subject is plotted. Particle variability is quantified as the standard deviation of per-trial particle likelihoods. The significant negative relationship indicates that lower particle variability is associated with longer response times. The mean and standard error is plotted. (B) Individual data points and the empirical distribution are plotted.

### Generalization to loss learning

To test whether our findings generalize beyond reward learning, we conducted an additional behavioral experiment, the turtle task, using the same prototypical 2×2 design that independently manipulated volatility and stochasticity. The turtle task is structurally identical to the sea lion task but implements loss avoidance rather than reward seeking, allowing us to test whether the dissociation between volatility and stochasticity holds across outcome valence. We recruited 30 participants for this experiment using Prolific.

Using the model-neutral blockwise HMM analysis, we first quantified learning rates for each block (Figure 7A). Replicating the results from the sea lion task, learning rates showed a significant positive main effect of true volatility (*t*(29) = +5.666, *P* < 0.001) and a significant negative main effect of true stochasticity (*t*(29) = −3.492, *P* = 0.002), with no significant interaction (*t*(29) = +0.704, *P* = 0.49; Supplementary Table 6). These results indicate that participants appropriately increased learning rate with volatility and decreased it with stochasticity, replicating our reward learning findings from the sea lion task.

**Figure 7.**
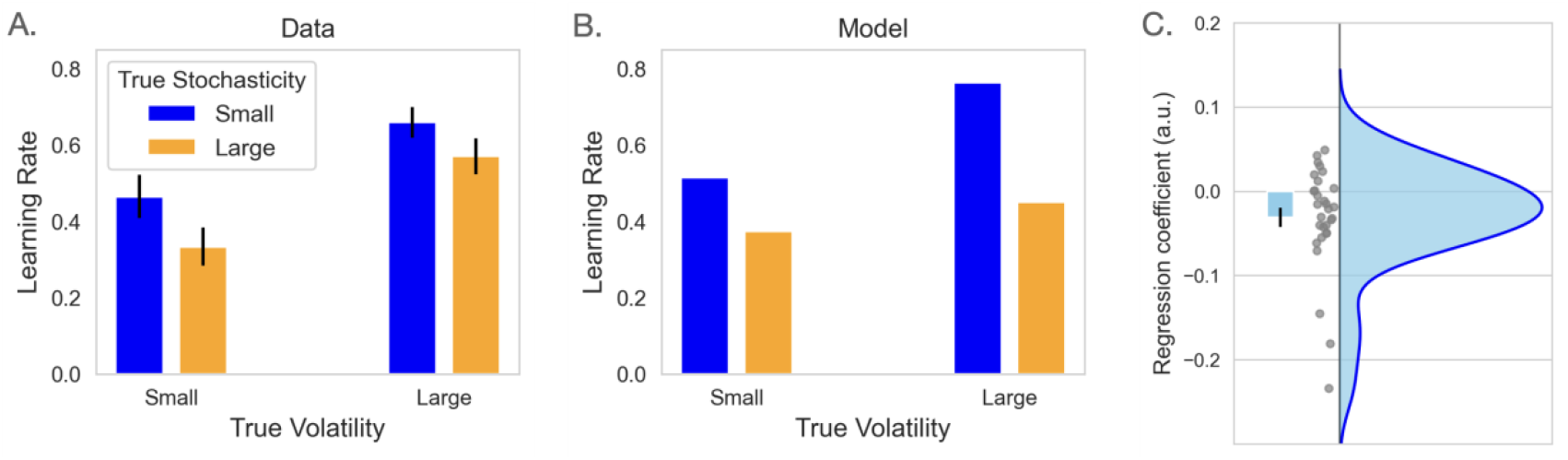
Generalization to an independent loss-learning task (turtle task). (A) Model-neutral analysis of an additional behavioral data with similar task setup. Mean learning rate per block is plotted (n = 30), which calculated by fitting the HMM model. The results show strong main effects of both factors: the learning rate increases with higher true volatility and decreases with higher true stochasticity. (B) Mean learning rate from fitting the PF-HMM model on subject data shows similar effects to model neutral analysis. (C) Linear regression results of participants’ response time (in seconds). The regression coefficient of particle likelihood variability (*PLV*) is plotted. The significant negative relationship indicates that lower particle variability is associated with longer response times. The mean and standard error, along with individual data points and the empirical distribution, are plotted.

We also repeated the response time analysis to test whether the PF-HMM predicts difficulty in inferring volatility versus stochasticity, as reflected in response times. Replicating the sea lion task results, response times were significantly larger when particle likelihood variability was low, indicating greater difficulty in identifying the source of uncertainty (*t*(29) = −2.799, *P* = 0.009; Figure 7C; Supplementary Table 7).

### Model comparison with alternative reversal-learning models

To assess whether the PF-HMM provides a better account of participants’ behavior than previous learning models, we compared it against two alternatives commonly used in reversal-learning tasks: the Pearce–Hall associability (PHA) model (Barnby et al., 2022; Li et al., 2011; Pearce & Hall, 1980) and the binary Hierarchical Gaussian Filter (HGF) (Iglesias et al., 2013; Mathys et al., 2011). All models were fitted to individual participants’ data by maximum likelihood estimation. Model evidence was calculated for each model using the Bayesian information criterion (Schwarz, 1978) and were used for random effects Bayesian model selection (Piray, 2026; Rigoux et al., 2014; Stephan et al., 2009). This procedure yields the model frequency, the expected proportion of participants best described by each model, and the protected exceedance probability, which quantifies the likelihood that a given model is the most frequent, while accounting for the possibility that observed differences arise by chance.

Bayesian model selection provided strong evidence in favor of the PF-HMM across both the sea lion and turtle tasks (Table 1). In the sea lion task, the PF-HMM achieved the highest model frequency (0.757) with a protected exceedance probability of 1.00, substantially exceeding both the PHA and HGF models. A similar pattern was observed in the turtle task, where the PF-HMM again dominated model comparison. Importantly, these alternative models fail both quantitatively and qualitatively: in addition to providing poorer fits to the data, they do not capture the model-neutral effects observed across datasets (Figures 5C and 7B), including the dissociable effects of volatility and stochasticity evident in the behavioral data (Supplementary Figure 5). In contrast, because the PF-HMM separately dissociates volatility from stochasticity, it accounts better for data, and it reproduces learning rate pattern observed across datasets.

**Table 1.**
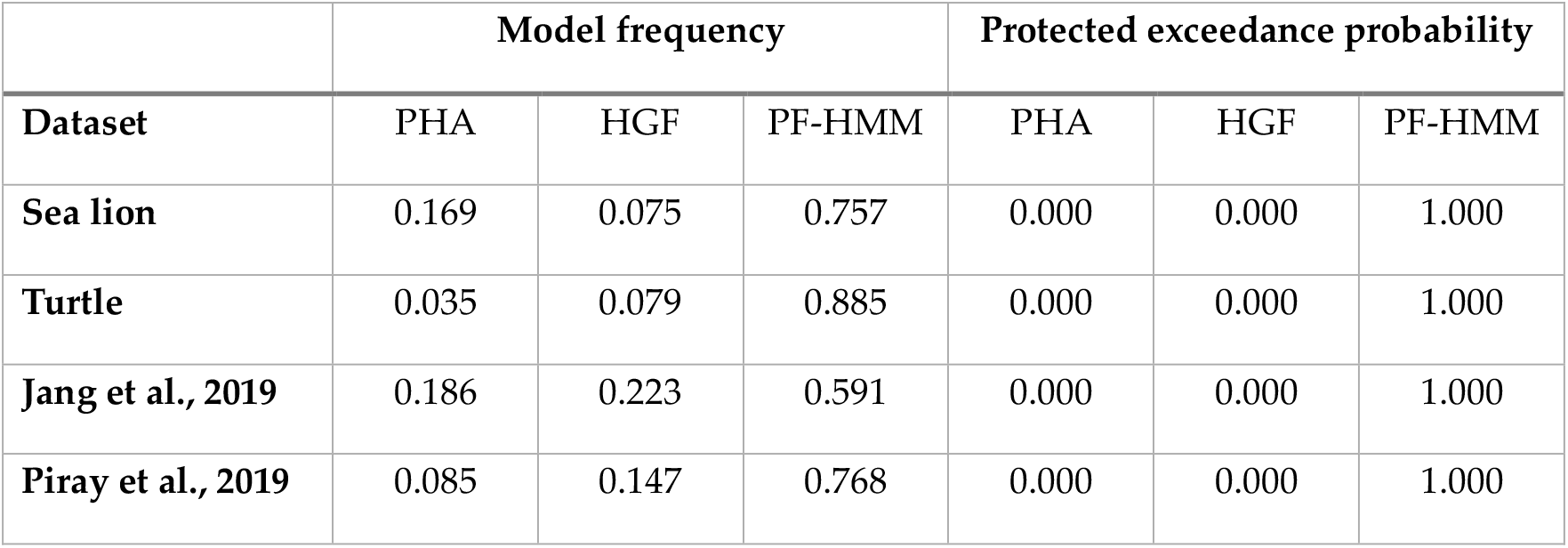
Bayesian model comparison results across four datasets.

|  | **Model frequency** |  |  | **Protected exceedance probability** |  |  |
| --- | --- | --- | --- | --- | --- | --- |
| **Dataset** | PHA | HGF | PF-HMM | PHA | HGF | PF-HMM |
| **Sea lion** | 0.169 | 0.075 | 0.757 | 0.000 | 0.000 | 1.000 |
| **Turtle** | 0.035 | 0.079 | 0.885 | 0.000 | 0.000 | 1.000 |
| **Jang et al., 2019** | 0.186 | 0.223 | 0.591 | 0.000 | 0.000 | 1.000 |
| **Piray et al., 2019** | 0.085 | 0.147 | 0.768 | 0.000 | 0.000 | 1.000 |

To assess the generality of these findings, we applied the same model fitting and Bayesian model selection procedures to two publicly available datasets involving probabilistic reversal learning. In the first dataset, 174 participants performed a reward-learning task in which they estimated the probability of reward based on binary feedback (Jang et al., 2019). On each trial, participants viewed an image and decided whether to accept or reject a gamble, with reward probability determined by the image category. Crucially, these contingencies shifted multiple times throughout the experiment, with each image category undergoing at least two reversals. The original study paired this learning task with a recognition memory test, administered entirely after the learning phase, to examine how reinforcement-learning dynamics shape episodic memory. Because the memory test did not intervene during learning, we can analyze the learning phase in isolation as an independent test of probabilistic learning under changing contingencies.

In the second dataset, 44 participants completed a probabilistic go/no-go task in which they learned to press or withhold a button in response to facial cues in order to obtain reward or avoid punishment (Piray, Ly, et al., 2019). The cues combined emotional content (happy or angry) with background color (grey or yellow for reward/punishment), and response–outcome contingencies for each cue were probabilistic and varied independently over time. The original studies examined socio-emotional modulation of learning and choice; here, we use the behavioral data from this task as a second independent test of learning under shifting contingencies. All learning models were augmented with a response model that included valence- and emotion-specific bias parameters to account for Pavlovian approach–avoidance tendencies, so differences in model comparison are attributable to learning dynamics alone.

Across both datasets, Bayesian model comparison again favored the PF-HMM (Table 1). The model achieved the highest global model frequency in the Jang et al. dataset (0.591) and the Piray et al. dataset (0.768), with protected exceedance probabilities of 1.000 in both cases.

## Discussion

Learning from binary outcomes is a fundamental challenge for organisms navigating uncertain environments. In such environments, observed outcomes provide only noisy, indirect information about an underlying hidden state that cannot be directly observed and may itself fluctuate over time. Despite the importance of understanding how organisms learn under these conditions, the literature has not yet addressed this problem from a comprehensive theoretical perspective. Instead, existing models have primarily relied on probabilistic algorithms designed for continuous outcomes (Browning et al., 2015; Iglesias et al., 2013; Mathys et al., 2011, 2014; Piray & Daw, 2020, 2021a), employing ad hoc approximations to handle binary data rather than systematically addressing the underlying statistical structure or considering how organisms might optimally learn from binary outcomes. This limitation is problematic even when stochasticity is fixed and known, as we have shown previously (Piray & Daw, 2020); see also (Ryali et al., 2018)). More importantly, it poses a fundamental challenge for understanding learning under conditions where both volatility and stochasticity are present, since continuous-based algorithms fail to distinguish between these distinct sources of noise.

We exploited the framework of probabilistic state space models, which serves as the underlying generative probabilistic graphical model for both the Kalman filter and HMM (Bishop, 2006; Ljung, 1998). While the Kalman filter is designed for continuous states and observations, the HMM framework is particularly suited to scenarios involving binary hidden states and binary outcomes (Ghahramani, 2001). Crucially, like the Kalman filter, the HMM supports exact inference, making it unnecessary to rely on Kalman-based approximate algorithms for handling binary outcomes. Additionally, while the Kalman filter recursively updates posterior statistics of the inferred hidden state—namely, its mean and variance—the HMM iteratively updates the posterior belief about the hidden state through a Bernoulli distribution, specifically estimating the probability that the state is 1 given outcomes.

Building on these parallels with the continuous case, we extended the concepts of volatility and stochasticity to the binary setting. Volatility refers to noise that disrupts the underlying binary hidden state over time, specifically the probability of the hidden state switching from one value to the other. In contrast, stochasticity reflects noise in the contingency between the hidden state and observed binary outcomes, specifically the probability that an outcome deviates from the true hidden state. From a statistical standpoint, optimal learning should adapt to these sources of uncertainty: Higher volatility requires faster learning because old information quickly becomes irrelevant. Conversely, higher stochasticity is expected to slow down learning, as each individual outcome is less informative, necessitating the integration of more outcomes to accurately infer the hidden state. We validated these predictions through simulation analyses using an optimal HMM with true values of volatility and stochasticity. As expected, optimal learning rates increased under high-volatility conditions and decreased under high-stochasticity conditions. To further test these predictions, we designed an experimental task employing a 2×2 factorial manipulation of volatility and stochasticity. Consistent with the optimal model’s behavior, human participants exhibited higher learning rates under high volatility and lower learning rates under high stochasticity.

We also developed a new model to capture learning in real-world situations where the true values of volatility and stochasticity are unknown and must be inferred from observed outcomes. To achieve this, we extended the HMM by incorporating a PF, a sequential Monte Carlo sampling method (Andrieu et al., 2010; Doucet et al., 2000). The new model, PF-HMM, allows volatility and stochasticity parameters to evolve dynamically on a trial-by-trial basis, reflecting real-time belief updates and closely mirroring participants’ incremental inference processes. The model integrates exact inference from the HMM with the PF’s per-trial estimates of volatility and stochasticity, enabling the model to continuously adapt to changing conditions while accurately distinguishing between different sources of noise. Simulation analyses demonstrated that the PF-HMM successfully inferred volatility and stochasticity, accurately attributing noise to its correct source. Importantly, this approach also allows us to predict two distinct patterns of impaired learning that arise from selective lesions to the model’s inference processes. Specifically, when the model’s ability to infer volatility is lesioned, it exhibits a maladaptive pattern characterized by a reduced learning rate, regardless of the true source of noise. Conversely, when the model’s ability to infer stochasticity is lesioned, it results in an excessive learning rate, indiscriminately adjusting to all forms of noise.

Previous experimental work has primarily focused on learning from binary outcomes under conditions of volatility (Brazil et al., 2017; Browning et al., 2015; Cole et al., 2020; de Berker et al., 2016; Deserno et al., 2020; Diaconescu et al., 2014, 2020; Hauser et al., 2014; Iglesias et al., 2013; Lawson et al., 2017; Piray, Ly, et al., 2019; Reed et al., 2020). While this line of research has been influential in both neuroscience and psychiatry, it has largely overlooked a more complex and challenging problem: dissociating volatility and stochasticity when both are unknown (Piray & Daw, 2021a, 2024). This distinction is crucial because volatility and stochasticity represent fundamentally different sources of, yet most previous studies have not systematically manipulated them experimentally. A notable exception is the recent work by Pulcu and Browning, who employed a 2×2 factorial design to manipulate both volatility and stochasticity, similar to our experimental design (Pulcu & Browning, 2025). However, unlike our findings, their study did not reveal any significant effects related to stochasticity (referred to as “noise” in their terminology). This discrepancy underscores the significance and robustness of our results, as well as the effectiveness of our experimental design. By successfully dissociating volatility and stochasticity, we established the theoretically expected opposing effects of these factors on learning rates: increased learning rates under high volatility and decreased learning rates under high stochasticity.

A related line of research has investigated learning from continuous observations under conditions of volatility and stochasticity, but until recently, it faced similar challenges in dissociating these distinct sources of uncertainty (Piray & Daw, 2021a). Our recent empirical work in that domain successfully addressed this issue by building on a well-established paradigm, demonstrating that learning rates increase with volatility and decrease with stochasticity (Piray & Daw, 2024). While both approaches employ PF models to dissociate volatility and stochasticity through dynamic tracking of these noise sources over time, the current model differs fundamentally at the level of inference. While both the previous model for continuous observations and the current model for binary observations use particle filter models to dissociate volatility and stochasticity by dynamically tracking these noise sources over time, they differ fundamentally in their inference process. The previous model relies on a Kalman filter for learning volatility and stochasticity based on continuous observations. In contrast, the model presented here is specifically designed for binary observations, employing a HMM to perform exact inference based on trial-by-trial estimates of volatility and stochasticity provided by the PF. This shift from Kalman filter-based inference to HMM-based inference is not merely a technical adjustment; it is a necessary adaptation to accurately capture the distinct statistical structure of binary outcomes. By integrating particle filtering with HMM, the current approach offers a novel and robust computational framework for dissociating volatility and stochasticity in binary learning problems, a challenge that has been unanswered by previous models focusing on continuous observations.

Beyond theoretical implications, accurately distinguishing volatility from stochasticity carries significant clinical relevance, particularly for understanding psychiatric disorders such as anxiety, schizophrenia, and impulse control disorders (Gagne et al., 2018; Horga & Abi-Dargham, 2019; Huang et al., 2017; Piray et al., 2014, 2017; Pulcu & Browning, 2019; Stephan et al., 2006; van Timmeren et al., 2023). These conditions often involve maladaptive responses to environmental uncertainty, where patients may exhibit either overly rigid or excessively flexible learning strategies (Browning et al., 2015; Lawson et al., 2017). Our model simulations, especially through targeted lesion experiments, provide valuable insights into how maladaptive estimation of volatility or stochasticity could drive clinically relevant maladaptive behaviors.

Our approach also highlights a critical issue overlooked by previous studies in computational psychiatry: the potential differences between learning from continuous and binary outcomes with respect to uncertainty-related parameters and their relevance to psychopathology. There are several key distinctions between these types of learning. Computationally, binary and continuous outcomes require fundamentally different algorithms for exact inference— specifically, Kalman filters for continuous outcomes and HMMs for binary outcomes. This distinction reflects underlying differences in how uncertainty is represented and processed within each framework. Psychopathologically, binary outcomes may be particularly salient in conditions such as depression, where individuals are prone to self-blame or heightened sensitivity to negative feedback (Abramson et al., 1978; Beck, 1970; Peterson et al., 1981; Zahn et al., 2015). Unlike continuous feedback, binary feedback offers clear-cut evaluations of success or failure, which may exacerbate maladaptive learning patterns and may partly explain why people learn differently from positive versus negative prediction errors (Frank et al., 2004; Piray, 2011; Pulcu & Browning, 2019). Specifically, when learning from negative feedback, individuals prone to self-blame may misattribute stochasticity (bad outcomes due to bad luck) to volatility (bad outcomes perceived as resulting from their own choices). This misattribution can reinforce negative biases by systematically interpreting perceived negative outcomes as evidence of environmental change, even when no such changes occur. Consequently, these individuals may develop maladaptive learning strategies that perpetuate their symptoms.

Although we focused on the problem of learning from binary outcomes under volatility and stochasticity, the dissociation of these two forms of uncertainty has potential implications for various cognitive processes that have not yet been extensively studied within this framework. One particularly relevant area is latent cause inference and structural learning, which involves determining underlying, categorical causes that govern observable patterns, such as stimuli, rewards, or punishments (Braun et al., 2010; Collins & Frank, 2013; Courville et al., 2004, 2006; Gershman et al., 2010; Lloyd & Leslie, 2013; Mahmoodi et al., 2024; Niv, 2019; Shin & Niv, 2021; Tomov et al., 2018). In these scenarios, the challenge lies in inferring which observations should be grouped together under a common cause and when a new cause should be inferred. The computational literature on latent cause inference has largely developed around probabilistic frameworks, such as the Chinese restaurant process (Gershman & Blei, 2012), which provides a principled way of inferring the emergence of novel causes based on observed data. These approaches have been influential in understanding how the brain constructs internal models of the environment, particularly when it encounters novel or changing structures (Gershman et al., 2010; Gershman & Blei, 2012; Lloyd & Leslie, 2013; Niv, 2019; Shin & Niv, 2021). Fundamentally, however, the problem involves clustering various observables in a way that accurately reflects their underlying causal structure (Courville et al., 2004, 2006). The ability to dissociate volatility and stochasticity is likely to play a critical role in this process. Specifically, if an agent infers similar volatility parameters for different observables, it suggests that those observables are likely changing together, potentially due to a shared underlying cause. This ability to track volatility separately from stochasticity provides a valuable signal for clustering observations appropriately, which is essential for constructing accurate causal models.

Another important issue that is theoretically relevant but have not been studied in the context of volatility and stochasticity is the issue of planning and decision making from a reinforcement learning perspective. Most previous work in this area has focused on prediction rather than optimal planning or decision-making. Even studies involving binary “choices” have primarily emphasized learning, as these tasks were essentially structured as prediction problems (Behrens et al., 2007, 2008; Browning et al., 2015). In fact, in these studies, the two choices were perfectly anti-correlated, and participants were explicitly informed of this structure. Consequently, participants’ learning processes could be assessed by measuring their ability to track changes in underlying contingencies, rather than by evaluating their capacity to make independent decisions aimed at maximizing reward, as is common in reinforcement learning (Sutton & Barto, 1998).

Extending the Bayesian generative processes underlying models of volatility and stochasticity to frameworks compatible with Markov decision processes (Sutton & Barto, 1998) commonly used in reinforcement learning presents substantial computational challenges. Prediction models typically focus on continuously updating beliefs about latent states, while planning and optimal control models must not only learn these parameters but also compute optimal sequences of actions that maximize future rewards over extended decision sequences (Sutton & Barto, 1998). Although some approaches have extended Kalman filtering to multi-step decision problems, resulting in Kalman-like temporal difference algorithms (Geist et al., 2009; Gershman, 2015), these methods fail to integrate the core optimal structure of either problem. Specifically, they do not combine the state space generative model characteristic of Kalman filtering with the optimal control framework essential to reinforcement learning. An alternative approach is the framework of planning as inference from control theory and machine learning (Kappen, 2005; Levine, 2018; Todorov, 2008, 2009; Toussaint & Storkey, 2006; Ziebart et al., 2008), which has recently influenced cognitive neuroscience research on planning, cognitive control, and cognitive map-making (Bazarjani & Piray, 2025; Botvinick & Toussaint, 2012; Piray & Daw, 2021b, 2025). This framework formulates optimal control as a Bayesian inference problem, providing a principled way to integrate uncertainty estimation with decision-making. By reframing planning as an inference problem, generative models of volatility and stochasticity can be incorporated directly into the generative model related to planning, rather than treating them as separate processes developed from different theoretical foundations and only superficially connected. This approach has the potential to unify prediction and control under a coherent computational framework. Overall, while significant progress has been made in understanding how agents learn to predict under conditions of volatility and stochasticity, extending these insights to reinforcement learning and planning remains a difficult and open problem. Achieving this integration requires reconciling models of uncertainty estimation with the demands of optimal control, including how agents select actions to achieve long-term goals rather than merely improving predictive accuracy.

While extending prediction models to decision-making introduces challenges related to long-term reward maximization and planning, a fundamental issue arises even in simpler multi-choice decision-making scenarios where sequential planning is unnecessary. Moving from prediction to decision-making inherently involves balancing exploration and exploitation (Amin et al., 2021; Daw et al., 2006; Song et al., 2019; Wilson et al., 2014). This trade-off requires deciding whether to exploit known options that provide reliable rewards or explore uncertain options that may yield better rewards in the future—a process strongly influenced by uncertainty estimates related to both volatility and stochasticity. Exploitation involves selecting actions based on existing knowledge to maximize immediate rewards, while exploration involves choosing actions that may be suboptimal in the short term but provide valuable information about the environment. In decision-making under uncertainty, agents must continually balance these competing demands to make adaptive choices. Multiple studies have shown that uncertainty estimates affect decision variability beyond what can be explained by learning and prediction alone (Findling et al., 2019, 2021; Lee et al., 2023). However, when both volatility and stochasticity are unknown and must be inferred from outcomes, distinguishing between prediction, behavioral variability, and information-driven exploration remains challenging.

## Conclusion

Inferring the true source of noise from binary outcomes is a fundamental challenge in real-world environments, especially under conditions of volatility and stochasticity where distinguishing between these sources is critical for adaptive learning. However, existing models designed for continuous data often rely on ad hoc approximations to handle binary outcomes, resulting in theoretical inconsistencies and limited generalization. To address this, we developed a novel framework, PF-HMM, which combines HMM with PF to infer volatility and stochasticity directly from binary outcomes. The model effectively dissociates these noise sources, providing a robust approach for understanding learning under uncertainty. Furthermore, lesion analyses using the PF-HMM reveal distinct patterns of maladaptive learning, offering potentially crucial insights into psychiatric conditions. Our experimental findings show that human participants adjust their learning rates appropriately, increasing them under high volatility and decreasing them under high stochasticity, consistent with model predictions. Participants’ response times are also slower when inferring the true source of noise is more difficult, as determined by the PF-HMM. Overall, this approach provides a principled framework for studying how organisms learn from binary outcomes and adapt to different sources of uncertainty in both healthy cognition and psychiatric conditions.

## Methods

### Experimental procedure

The study was approved by the Institutional Review Board at the University of Southern California (“Learning and decision making under uncertainty”; Protocol Number: UP-23-00359). Participants were recruited online via the Prolific platform and provided informed consent prior to beginning the experiment. This study was not preregistered. The task was implemented in JavaScript and jsPsych (Leeuw et al., 2023) and deployed using the NivTurk platform (Zorowitz & Bennett, 2022).

Participants first read instructions, completed step-by-step practice trials to familiarize themselves with the task, and then completed a comprehension quiz to ensure understanding. Those who failed the quiz twice were not allowed to proceed. A total of 73 participants were recruited for the sea lion task. The sample size was chosen based on pilot data from a separate sample, which suggested that it would yield at least 0.8 statistical power. A total of 30 participants were recruited for the turtle task, based on a power analysis using the sea lion task data. All participants passed the comprehension check and met quality control criteria, including consistent engagement with the task (i.e., not leaving the interface idle for extended periods). After completing the main behavioral task, participants also completed an additional task (not analyzed here) and the demographic questionnaire. Data from none of the participants were excluded from the analyses.

### Blockwise HMM

To obtain model-neutral estimates of learning rate, we fitted a binary Hidden Markov Model (HMM) separately to each block of each participant’s data. The full generative formulation and filtering equations are provided in the Appendix A.3. Briefly, the HMM assumes a latent binary state that evolves according to a first-order Markov process governed by volatility *v*, and generates outcomes through a stochasticity parameter *s*.

For each block, two parameters were fitted: *v* volatility (probability of hidden state switching) and *s* stochasticity (probability that the observed outcome deviates from the hidden state). Given these parameters, the HMM recursively computes the predicted probability that the hidden state equals 1, denoted as *r*_*t*_. These predicted probabilities reflect belief updating under the assumed volatility and stochasticity levels.

### Response model

Choices were modeled using a response function that maps model-derived predicted probabilities to observed responses. While the HMM provides the predicted belief about outcome probability, actual choices may also reflect motor-related perseveration. To account for this, predicted probabilities were transformed into log-odds space and augmented by a stickiness parameter *ρ*, capturing the tendency to repeat or switch the previous choice. This response-level parameter operates independently of belief updating and latent state inference.

Let *p*_*t*_ denote the final predicted probability of choosing option 1 after applying the response model. The same response model structure was applied across all fitted models to ensure comparability of likelihood evaluation.

### Model fitting procedure for blockwise HMM

Parameters were estimated separately for each block using maximum a posteriori (MAP) estimation. Raw parameters were optimized in unconstrained space and transformed via a half-sigmoid function 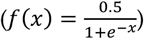 to enforce valid ranges. A zero-mean Gaussian prior with fixed variance was placed on each parameter in the unconstrained space. MAP estimation was implemented using the CBM package (Piray, Dezfouli, et al., 2019), which employs a Laplace approximation around the posterior mode and provides an estimate of model evidence. We used a wide variance (100) to ensure that parameters could vary over a wide range.

For each trial, the HMM filtering equations were used to compute *r*_*t*_, which was then passed through the response model to obtain *p*_*t*_. The log-likelihood of observed choices was computed as:

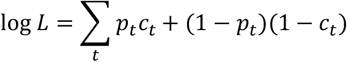

where *c*_*t*_ is the choice (0 or 1) on trial *t*. Trials with missing responses were excluded from the likelihood computation. Blockwise log-likelihoods were summed across blocks for subject-level model fitting.

We also performed a recovery analysis to assess how well model parameters can be recovered. We generated N=100 synthetic datasets by sampling model parameters randomly. The synthetic datasets were subsequently fitted using the same model-fitting procedure employed for empirical data. This analysis revealed that model parameters are highly recoverable (see Supplementary Table 1 and Supplementary Fig. 3).

### Model fitting procedure for PF-HMM

Fitting the PF-HMM poses substantial challenges due to the stochastic nature of particle filtering and the lack of differentiability of the likelihood with respect to model parameters. We therefore employed a gradient-free optimization approach based on Bayesian optimization, implemented using MATLAB’s bayesopt routine. Model parameters were fit separately for each participant by maximizing the log-likelihood of observed behavioral data under the PF-HMM. All reported PF-HMM fits include a response-level perseveration parameter, which accounts for choice repetition effects independently of latent state inference (see Response model below). For completeness, we report a comparison between PF-HMM fits with and without the perseveration parameter in the Supplementary Materials (Supplementary Table 8), demonstrating that inclusion of the response model improves model fit without altering the qualitative learning-rate effects.

Our approach is similar to our recent work for fitting PF models to human data (Piray & Daw, 2024). Specifically, for a given parameter setting, model likelihood was evaluated by running the particle filter multiple times (10 runs per evaluation) to account for stochastic variability in the inference process. The objective function minimized by the optimizer was defined as the mean negative log-likelihood across runs, augmented by a penalty proportional to the standard error of the mean across runs. This regularized objective favors parameter estimates that both achieve high likelihood and yield stable estimates across repeated stochastic simulations.

Bayesian optimization was initialized with multiple seed points and iteratively evaluated candidate parameter settings using an expected-improvement acquisition function. For each participant, the best-fitting parameters were selected based on the optimized objective. Latent trajectories—including inferred beliefs about the hidden state, volatility, and stochasticity— were obtained by averaging model outputs across repeated particle filter runs using the fitted parameters. Learning rates were computed post hoc as descriptive summaries by regressing belief updates on prediction errors derived from the model’s latent state estimates (see below).

To assess parameter recoverability, we conducted a recovery analysis in which synthetic datasets were generated by simulating the PF-HMM using randomly sampled parameter values. The same fitting procedure was then applied to these synthetic datasets. This analysis demonstrated high recoverability for all model parameters (Supplementary Table 4 and Supplementary Fig. 4).

A recovery analysis was conducted for the PF-HMM with perseveration. Again, *N* = 100 synthetic datasets were generated. True parameters were sampled from the same ranges used during model fitting, including the diffusion parameters governing volatility and stochasticity and the global perseveration parameter. For each simulated subject, belief trajectories were generated using the full PF-HMM forward model with the same particle-filter configuration as used in fitting (10,000 particles, systematic resampling, identical initialization). To reduce stochastic variability, belief trajectories were averaged across multiple forward passes. Choices were then generated sequentially using the same response model.

### Learning rate

For HMM and the PF-HMM models, the learning rate was defined as the ratio of update to prediction error. The belief update *Δr*_*t*_ measures how much the posterior belief, *r*_*t*_, changes after observing the outcome:

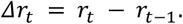

The prediction error *δ*_*t*_ quantifies the discrepancy between the observed outcome *o*_*t*_ and the model’s prediction on trial *t*:

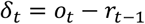

The trial-by-trial learning rate is defined as the ratio of the belief update to the prediction error, capturing how strongly new information influences belief revision. However, in practice, we quantified learning rates at the block level by regressing belief updates onto prediction error across trials within each block:

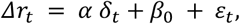

where *α* is the regression coefficient related to prediction errors (i.e., the learning rate for that block), *β*_0_ is learning-independent block effect, and *ε*_*t*_ is noise.

### Response time analysis

To examine whether computational signatures of noise-source attribution are reflected in behavior beyond choice, we conducted a trial-by-trial regression analysis of response times (RTs). The analysis tested the hypothesis that responses would be slower when identifying the source of noise is more difficult, reflected as low particle variability in the PF-HMM.

For each trial, model predictions were generated using the median fitted parameter values across participants. From these trajectories, we extracted:

- Particle likelihood variability (*PLV*_*t*_): the standard deviation of per-particle likelihoods at trial *t*.
- Average per particle likelihood (*PLM*_*t*_): the average per-particle likelihood at trial *t*.
- Estimated volatility (*V*_*t*_): the PF-HMM estimate of volatility at trial *t*.
- Estimated stochasticity (*S*_*t*_): the PF-HMM estimate of stochasticity at trial *t*.
- Accuracy (*Acc*_*t*_): a binary indicator (1 if the participant’s choice matched the outcome; 0 otherwise).

Response times were extracted from trial *t*, while model-derived regressors were aligned to trial *t* − 1 to ensure proper temporal ordering between predictors and behavioral response. Outlier response times were identified and removed using standard outlier detection procedures.

For each participant, we fit the following linear regression model:

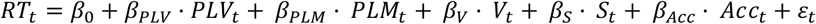

Here, *RT*_*t*_ denotes the response time on trial *t*, and *ε*_*t*_ is noise. The regression model was fit separately for each participant separately, yielding subject-specific regression coefficients. To assess group-level effects, we performed one-sample *t*-tests on the regression coefficients across participants. Mean effects, standard errors of the mean (SEM), *t*-values, and *p*-values were computed for each regressor.

## Code and data availability

Analyses were conducted in MATLAB (R2023b) and Python (3.11.5). All experimental data and analysis code are publicly available online at https://github.com/piraylab/PF-HMM. The manuscript is available on *bioRXiv*.

## Acknowledgements

This work was supported by grants R21MH134217 from the National Institute of Mental Health. We thank Nathaniel Daw for helpful discussions. The authors declare no conflicts of interest.

## Appendix A

### A.1 State space models

We begin by describing state space models (SSMs), which provide a general probabilistic framework for modeling dynamical systems in which a sequence of observations is generated from an underlying sequence of latent states. Let 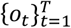 denote the observed outcomes and 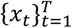 denote the corresponding latent states. In general, SSMs might have optional inputs, and might also have

Note that the term state space model is used broadly to refer to any recurrent process with a latent state. In general, SSMs may be noisy or noiseless, may be driven by external inputs, and may involve latent states that are one-or multi-dimensional. Here, we focus on noisy SSMs with a one-dimensional latent state and observation.

SSMs assume the following conditional independence structure:

1. First-order Markov process of the latent state:

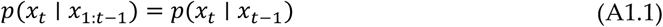
2. Conditional independence of observations given the latent state:

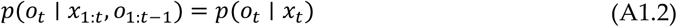

The primary inference goal (filtering) in an SSM is to compute the posterior over the current latent state given observations up to time *t*, i.e. *p*(*x*_*t*_ ∣ *o*_1:*t*_). Importantly, this posterior can be computed recursively, regardless of the specific functional forms of the transition and

observation models. This typically is done in two steps:

- Prediction: Given the posterior at time *t* − 1, *p*(*x*_*t*−1_ ∣ *o*_1:*t*−1_), the predicted distribution at time *t* (prior to observing *o*_*t*_) is obtained by marginalizing over the previous state:

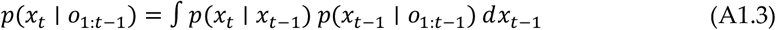
- Update: After observing *o*_*t*_, the estimate is adjusted by actual measurement using Bayes’ rule, giving rise to the posterior given all observations:

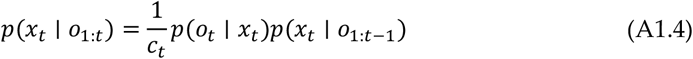

where *c*_*t*_ is the normalization constant with respect to *x*_*t*_:

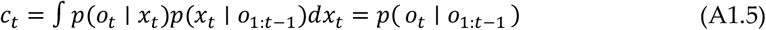

Equations (A1.3 – 1.5) hold for any model satisfying the conditional independence structure above, irrespective of the functional forms of *p*(*x*_*t*_ ∣ *x*_*t*−1_)and *p*(*o*_*t*_ ∣ *x*_*t*_). However, the tractability of the integral in Eq. (A1.3) depends on those forms. Two important special cases admit exact recursive solutions. In linear–Gaussian SSMs, both the transition and observation models are Gaussian, and the filtering distribution *p*(*x*_*t*_ ∣ *o*_1:*t*_) remains Gaussian at every step. Exact inference in this setting is provided by the Kalman filter. Although the Kalman filter applies more generally to multidimensional systems, we briefly review the one-dimensional case below. In discrete-state SSMs, where latent states take values in a finite set, the filtering distribution is categorical, and the recursion reduces to finite summation rather than integration. Exact inference in this setting is provided by the hidden Markov model (HMM) filtering equations. In the present work, we focus on the binary system, in which *x*_*t*_ ∈ {0,1} and *o*_*t*_ ∈ {0,1}.

### A.2 Kalman filter

The Kalman filter arises as a special case of the SSM under linear transition dynamics and Gaussian noise. It provides an exact recursive solution when both the hidden state and the observations are continuous-valued, and all noise sources are Gaussian:

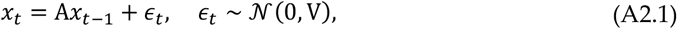

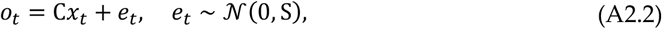

where *x*_*t*_ denotes the hidden state, *o*_*t*_ denotes the observation, and *ϵ*_*t*_ and *e*_*t*_ are independent Gaussian noise whose covariances is given by V and S, respectively. We focus on the special scalar case with A = C = 1, V = *v* (volatility), and S = *s* (stochasticity). In probabilistic form, these become:

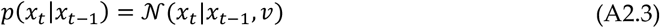

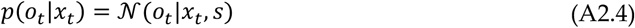

Under these assumptions, it can be shown that the posterior remains Gaussian at every time step:

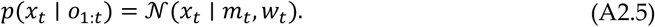

Thus, inference reduces to recursively updating the posterior mean *m*_*t*_ and variance *w*_*t*_. Specifically, inference proceeds in two steps: prediction and update, corresponding to Eqs. (A1.3 – 1.4) in the general SSM formulation. The predicted mean and covariance prior to observing *o*_*t*_, 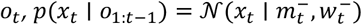, are obtained by propagating the previous posterior through the linear dynamics. It can be shown that:

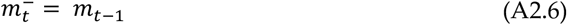

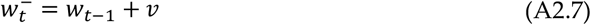

After observing *o*_*t*_, the posterior is given by Equation A1.4. Defining the Kalman gain as:

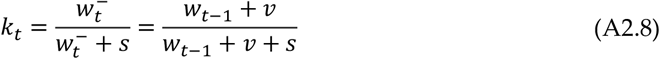

the posterior mean and variance are then given by:

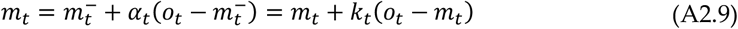

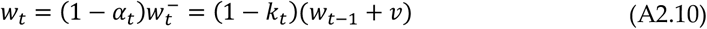

Thus, the Kalman gain *k*_*t*_ determines how strongly new observations influence the posterior estimate. It increases with *v* (volatility) and decreases with *s* stochasticity. In this setting, learning rate, defined as update divided by prediction error, (*m*_*t*_ − *m*_*t*−1_)/(*o*_*t*_ − *m*_*t*−1_), is equivalent to the Kalman gain.

### A.3 Hidden Markov model

We consider a binary HMM, in which both hidden states and observations take values of either 0 or 1. The hidden state evolves according to a binary diffusion process governed by volatility parameter *v*. Specifically,

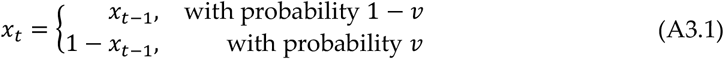

where *v* ∈ [0, 0.5] denotes the probability that the hidden state switches on trial *t*. When *v* = 0, the state is perfectly stable; when *v* = 0.5, the state is completely noisy. The corresponding conditional distribution of *x*_*t*_ can be written compactly as:

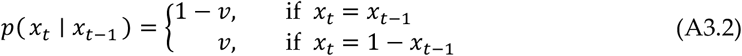

Observations are generated by corrupting the hidden state with observation noise governed by the stochasticity parameter *s*:

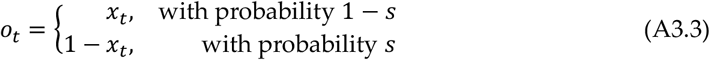

where *s* ∈ [0, 0.5] is the stochasticity parameter indicating how noisy the observation process is. Larger values of *s* indicate noisier observations, with *s* = 0.5 indicating that the observation is equally likely to match or differ from the latent state *x*_*t*_. In contrast, when *s* = 0, the observation matches the state. Thus, the conditional distribution of *o*_*t*_ can be written as:

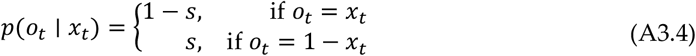

Under these assumptions, it can be shown that the posterior at every time step takes the form of a Bernoulli distribution. In other words, for binary SSMs, the Bernoulli distribution plays the role that the Gaussian plays in linear-Gaussian SSMs. Recall that for a binary random variable *z* ∈ {0,1}, the Bernoulli distribution with parameter *μ* ∈ [0, 1] is given by:

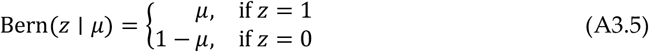

Thus, parameter *μ* represents probability of *z* = 1.

Now suppose the posterior belief at time *t* − 1 is Bernoulli with parameter *r*_*t*−1_,

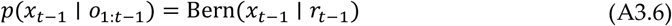

where *r*_*t*−1_ = *p*(*x*_*t*−1_ = 1 ∣ *o*_1:*t*−1_) represents the probability that the hidden state equals 1 given all outcomes up to and including time *t* − 1. We now show that the posterior at time *t* is also Bernoulli.

Since the binary HMM satisfies the conditional independence structure of the general state space model (Section A.1), the recursive update equations (A1.3–1.4) apply directly. The prediction step computes the belief about the current state before observing *o*_*t*_, that is, *p*(*x*_*t*_ = 1 ∣ *o*_1:*t*−1_). Using the general SSM recursion (Eq. A1.3) with integration replaced by summation, the predicted distribution is obtained by marginalizing over the previous hidden state:

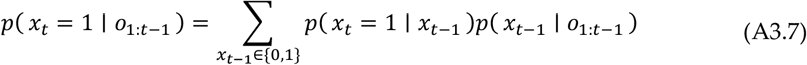

Substituting the transition model (Eq. A3.2) and definition of *r*_*t*−1_, we obtain:

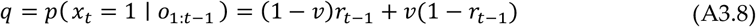

This yields Eq. (5) in the main text. Thus, *q* represents belief about that *x*_*t*_ = 1 before incorporating the current observation. Similarly, we have *p*(*x*_*t*_ = 0 ∣ *o*_1:*t*−1_) = 1 − *q*, and consequently *p*(*x*_*t*_ ∣ *o*_1:*t*−1_) = Bern(*x*_*t*_ ∣ *q*). While not needed for the HMM update, it is useful for the next section to also obtain *p*(*o*_*t*_|*o*_1:*t*−1_) before observing *o*_*t*_:

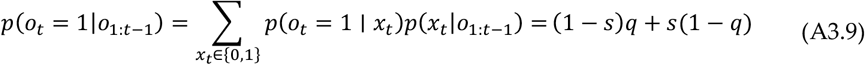

Next, we use Eq (A1.4) to calculate the posterior after observing *o*_*t*_. First, the likelihood of the observed outcome *o*_*t*_ under the assumption that *x*_*t*_ = 1, can be written as:

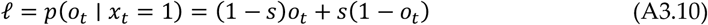

This corresponds to Eq. (6) in the main text. Note that *l* depends only on the stochasticity parameter *s* and the observed outcome *o*_*t*_. Similarly, from the observation model, we obtain *p*(*o*_*t*_ ∣ *x*_*t*_ = 0) = *so*_*t*_ + (1 − *s*)(1 − *o*_*t*_) = 1 − *l*.

Second, applying Eq. A1.4 yields the posterior belief after observing the current outcome:

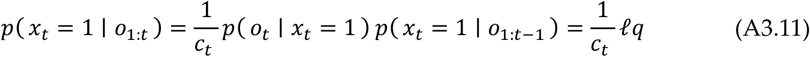

and similarly:

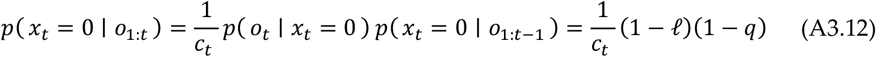

Since, *p*(*x*_*t*_ ∣ *o*_1:*t*_) is a probability distribution, it must sum to one, which yields the normalizing constant, *c*_*t*_:

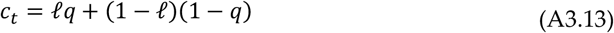

Thus, the Bernoulli form is preserved under recursive Bayesian updating, *p*(*x*_*t*_ ∣ *o*_1:*t*_) = Bern(*x*_*t*_ ∣ *r*_*t*_), with

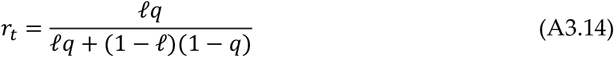

This corresponds to Eq. (7) in the main text.

### A.4 Particle filter-HMM

Now we extend the binary SSM by treating both volatility and stochasticity as dynamically changing random variables. Since their values are bounded between 0 and 0.5, we model these processes using the Beta distribution. Specifically, we define a reparametrized version of the Beta distribution, ℬ(*z*|*μ*, σ), as follows:

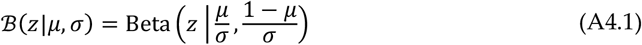

Here, Beta denotes the standard Beta distribution. This reparameterization ensures that the expected value of *z* is *μ*, while σ controls the dispersion of the distribution. Using this formulation, we define the volatility and stochasticity processes as:

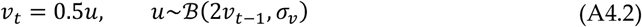

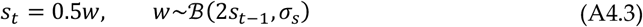

where σ_*v*_ and σ_*s*_ determine the stepwise diffusion for volatility and stochasticity, respectively. Importantly, the conditional expected values of *v*_*t*_ and *s*_*t*_ are *v*_*t*−1_ and *s*_*t*−1_, respectively, thereby maintaining the martingale property of the process.

The hidden state and outcomes are generated through the same generative process underlying the HMM on each trial:

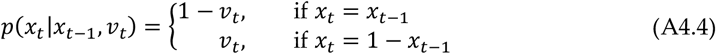

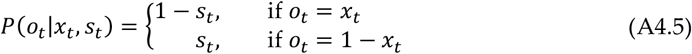

Since exact inference is no longer tractable, we use Rao-Blackwellized particle filtering (Doucet et al., 2000; Doucet & Johansen, 2011): each particle *i* maintains estimates 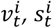, and 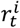, updated in three steps per trial.

First, for each particle *i*, volatility and stochasticity transition forward in time according to their generative dynamics. This gives rise to sampled values 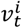 and 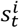 Specifically, we sample these from the generative distributions defined in Eqs. (A4.2–A4.3):

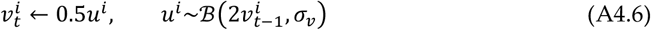

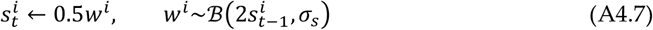

The second step updates particle weights based on outcome likelihood:

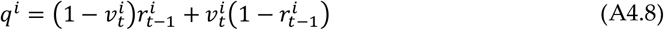

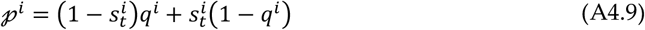

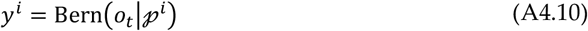

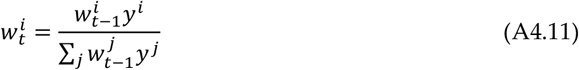

where 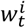 represents particle *i*’s weight on trial *t*. The intermediate variable *q*^*i*^ represents the probability of the current state being 1 conditional on 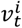 and 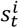 before observing the current outcome. The intermediate variable 𝓅^*i*^ is the predictive probability of the outcome *o*_*t*_ being 1, conditional on 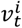 and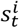. As is customary in PF methods, systematic resampling is applied during this step if the effective number of particles drops below a certain threshold.

The third step applies the HMM to integrate the new outcome with previous estimates. For each particle, we update 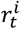 using HMM update equations.

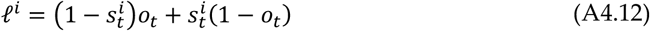

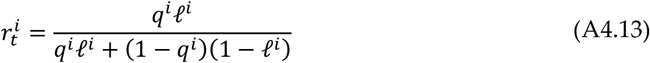

The expected value for all random variables at trial *t* is computed as the weighted average across all particles, with *w*^*i*^ serving as the weighting coefficients.

## Appendix B

In the main text, we noted that binary models built on the Kalman filter exhibit a problematic relationship with the learning rate. Here, we formalize this claim. This argument appeared in our earlier work (Piray & Daw, 2020); here, we present it more slightly more generally and include it for completeness. Specifically, we show that a class of generative models based on sigmoid transformations of linear–Gaussian state-space models, and the resulting Kalman-filter-based algorithms, such as hierarchical Gaussian filters (HGF) and the volatile Kalman filter, do not yield the correct relationship between learning rate and volatility.

These models assume the following conditional dependencies:

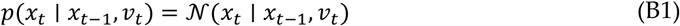

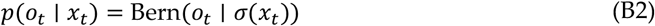

where σ(*x*) = 1/(1 + exp (−*x*))is the sigmoid function. Here, *v*_*t*_ is volatility, which is itself dynamically changing. While different models make different assumptions about the dynamics of *v*_*t*_, our analysis is independent of these assumptions. In fact, our argument holds even if volatility is fixed, i.e., *v*_*t*_ = *v*. Under these models, the posterior over *x*_*t*_ is approximated as Gaussian. Its mean, *m*_*t*_, takes the form:

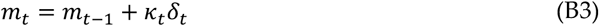

where κ_*t*_ is the modified Kalman gain, which typically depends on inferred volatility, and *δ*_*t*_ = *o*_*t*_ − σ(*m*_*t*−1_) is the prediction error. Since *o*_*t*_ ∈ {0,1} and the sigmoid is bounded between 0 and 1, it follows that −1 < *δ*_*t*_ < 1.

We now show that the learning rate is directly contaminated by large prediction errors, regardless of κ_*t*_ and regardless of whether the prediction error is driven by volatility. The learning rate is defined as:

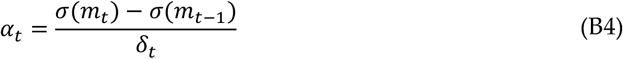

Substituting the update equation for *m*_*t*_:

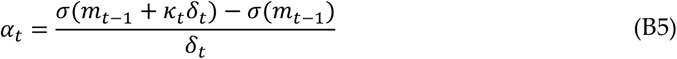

The key point is that the numerator of α_*t*_ depends on second-order effects of *δ*_*t*_. To see this, we expand σ(*m*_*t*−1_ + κ_*t*_*δ*_*t*_) using a Taylor series around *m*_*t*−1_:

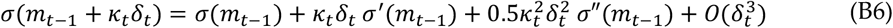

where σ^′^and σ″ denote the first and second derivatives of the sigmoid, and 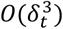 represents terms of order 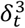 and higher (that are small given that |*δ*_*t*_| < 1. Substituting into the expression for α_*t*_:

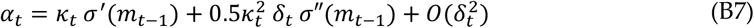

For the sigmoid function:

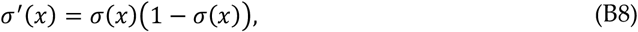

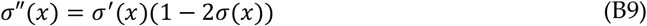

Note that σ^′^(*x*) ∈ [0,0.25] and (1 − 2σ(*x*)) ∈ [−1,1]. Substituting:

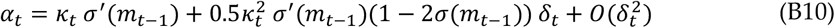

We are interested in the dependence of α_*t*_ on the prediction error *δ*_*t*_. The key term is (1 − 2σ(*m*_*t*−1_)) *δ*_*t*_, which is large in magnitude whenever |*δ*_*t*_| is large, regardless of volatility and its effects on κ_*t*_. In particular, there are two cases corresponding to large prediction errors:

- *o*_*t*_ = 1 and *m*_*t*−1_ is large and negative: both (1 − 2σ(*m*_*t*−1_)) and *δ*_*t*_ are positive and large.
- *o*_*t*_ = 0 and *m*_*t*−1_ is large and positive: both (1 − 2σ(*m*_*t*−1_)) and *δ*_*t*_ are negative and large.

Thus, in both cases, α_*t*_ is inflated by |*δ*_*t*_|. In other words, the learning rate is inherently contaminated by the magnitude of the prediction error, regardless of the source of noise, and independent of volatility estimate affecting κ_*t*_.

