## Supplementary Information for "Inferring the causes of noise from binary outcomes: A normative theory of learning under uncertainty"

### Supplementary Methods

#### Hierarchical Gaussian Filter

We implemented a three-level Hierarchical Gaussian Filter (HGF) following the formulation of Mathys et al. (2011). The HGF resembles a state-space model in which latent states evolve as coupled Gaussian random walks. The generative process is defined as follows:

$$x_3^{(t)} \sim \mathcal{N}(x_3^{(t)} | x_3^{(t-1)}, \nu),$$
$$x_2^{(t)} \sim \mathcal{N}(x_2^{(t)} | x_2^{(t-1)}, \exp(\kappa x_3^{(t)} + \omega)),$$
$$x_1^{(t)} = x_2^{(t)},$$
$$o_t \sim \text{Bern}\left(\sigma(x_1^{(t)})\right),$$

where $x_1$, $x_2$, and $x_3$ are dynamic random variables, $o_t$ is the observed outcome, and $\sigma(x) = \frac{1}{1+\exp(-x)}$ is the sigmoid function. $\nu$, $\kappa$, and $\omega$ are model parameters. Thus, $\nu$ governs changes in $x_3$, which in turn determines the variance of belief updates for $x_2$. Consequently, the volatility that effectively influences $x_2$ varies as an exponential function of $x_3$ parameterized by the $\kappa$ and $\omega$ parameters. Inference in the HGF is performed using a variational approximation under the Laplace assumption, yielding closed-form update equations for posterior means and variances at each level (Mathys et al., 2011). In particular, under these assumptions, the posterior over $x_2$ takes a Gaussian form, in which its mean, $\mu_t$, takes a Kalman-form update:

$$\mu_t = \mu_{t-1} + \kappa_t(o_t - \sigma(\mu_{t-1})),$$

where $\kappa_t$ is the modified Kalman gain that depends on volatility. This update equation is needed for the proof presented in Appendix B. We refer readers to the original papers for the full mathematical details.

For model fitting, the three parameters were fitted along task-specific response parameters (e.g., perseveration). Similar to fitting PF-HMM, we performed maximum likelihood to estimate model parameters. Parameters were estimated independently for each participant using constrained nonlinear optimization (MATLAB fmincon). Learning rate was computed per block using the same regression analysis presented in the main text.

#### Pearce–Hall Associability Model

To compare the PF-HMM against a classical surprise-driven learning framework, we implemented a Pearce–Hall associability (PHA) model adapted for binary outcomes. In this model, learning is governed by trial-wise prediction errors and a dynamically evolving associability parameter that determines how strongly new outcomes influence belief updating.

The model maintains a latent belief state $V_t \in [0, 1]$, representing the learner's estimate of the probability that the outcome equals 1 prior to observing the outcome at trial $t$. On each trial, the predicted probability of outcome 1 is simply given by $V_t$. After observing the binary outcome $o_t \in \{0, 1\}$, a prediction error is computed as:

$$\delta_t = o_t - V_t.$$

Beliefs are updated according to a delta rule:

$$V_{t+1} = V_t + \alpha_t \delta_t,$$

where $\alpha_t \in [0, 1]$is the associability parameter, which functions as a dynamic learning rate. Importantly, in the Pearce–Hall framework associability evolves as a function of unsigned prediction error, reflecting the principle that surprising outcomes increase attention to subsequent stimuli. Specifically, associability is updated according to:

$$\alpha_{t+1} = (1 - \kappa)\alpha_t + \kappa \mid \delta_t \mid,$$

where $\kappa \in (0, 1)$ controls the rate at which associability adapts to recent surprise. Larger values of $\kappa$ result in more rapid increases in learning rate following unexpected outcomes, whereas smaller values lead to slower adaptation. Initial conditions were set to $V_0 = 0.5$ and $\alpha_0 = 1$, corresponding to an unbiased initial belief and maximal initial associability. Parameters were estimated independently for each participant using constrained nonlinear optimization (MATLAB fmincon). The log-likelihood of observed choices was computed from predicted choice probabilities, excluding trials with missing responses. Learning rate was computed per block using the same regression analysis presented in the main text.

| | 25% quantile | Median | 75% quantile |
|---|---|---|---|
| $v_1$ | -0.030 | 0.004 | 0.053 |
| $v_2$ | -0.013 | 0.021 | 0.131 |
| $v_3$ | -0.062 | -0.001 | 0.059 |
| $v_4$ | -0.034 | 0.013 | 0.089 |
| $s_1$ | -0.016 | 0.009 | 0.038 |
| $s_2$ | -0.022 | 0.008 | 0.039 |
| $s_3$ | -0.013 | 0.009 | 0.044 |
| $s_4$ | -0.030 | 0.011 | 0.054 |
| $\rho$ | -0.103 | 0.102 | 0.315 |

Supplementary Table 1. Recovery analysis of the HMM with perseveration. The table shows summary statistics of errors between true (simulated) parameters and those recovered by the model. The 25%, 50% (Median), and 75% quantiles of the error distributions are reported. Both volatility ($v_1$ to $v_4$) and stochasticity parameters ($s_1$ to $s_4$) show median recovery errors close to zero, indicating effective recovery of parameters with no systematic bias.

| | Sea lion task | | Turtle task | |
|---|---|---|---|---|
| | Mean | STD | Mean | STD |
| $v_1$ | 0.230 | 0.176 | 0.267 | 0.179 |
| $v_2$ | 0.306 | 0.166 | 0.301 | 0.184 |
| $v_3$ | 0.199 | 0.186 | 0.159 | 0.164 |
| $v_4$ | 0.303 | 0.162 | 0.298 | 0.202 |
| $s_1$ | 0.214 | 0.147 | 0.246 | 0.143 |
| $s_2$ | 0.149 | 0.125 | 0.149 | 0.103 |
| $s_3$ | 0.260 | 0.165 | 0.283 | 0.145 |
| $s_4$ | 0.227 | 0.153 | 0.208 | 0.126 |
| $\rho$ | 0.668 | 1.104 | 0.622 | 0.774 |

Supplementary Table 2. Summary statistics of HMM with perseveration fitted parameters. The three columns show the mean and standard deviation for the fitted model parameters. $v_1$ and $s_1$ are volatility and stochasticity parameters, respectively, for block 1 (small true stochasticity and small true volatility); $v_2$ and $s_2$ for block 2 (small true stochasticity and large true volatility); $v_3$ and $s_3$ for block 3 (large true stochasticity and small true volatility); and $v_4$ and $s_4$ for block 4 (large true stochasticity and large true volatility).

| | $\delta$ | $\delta \times S$ | $\delta \times V$ | $\delta \times S \times V$ | $S$ | $V$ | $S \times V$ | $I$ |
|---|---|---|---|---|---|---|---|---|
| Coeff. | 1.032 | -0.101 | 0.145 | -0.024 | 0.003 | 0.002 | -0.009 | -0.007 |
| S.E.M. | 0.055 | 0.023 | 0.028 | 0.020 | 0.001 | 0.001 | 0.002 | 0.001 |
| t-value | 18.764 | -4.302 | 5.286 | -1.242 | 2.113 | 2.343 | -5.322 | -6.739 |
| p-value | <0.001 | <0.001 | <0.001 | 0.218 | 0.038 | 0.022 | <0.001 | <0.001 |

Supplementary Table 3. Fitted blockwise HMM to sea lion task data (n=73). This table presents the regression results for all regressors. The columns list each regressor with corresponding coefficient estimates (Coeff.), standard errors of the mean (S.E.M.), t-values, and p-values form two-sided t-tests. The term $\delta$ is defined as the observation minus previous belief ($\delta_t = o_t - r_{t-1}$). The factors $S$ and $V$ refer to the true stochasticity and true volatility, respectively, while $I$ represents the intercept term. Interaction terms (e.g., $\delta \times V$, $\delta \times S$, etc.) capture interaction effects of $\delta$ with the true environment parameters.

| | 25% quantile | Median | 75% quantile |
|---|---|---|---|
| $\sigma_s$ | -0.051 | 0.003 | 0.067 |
| $\sigma_v$ | -0.048 | 0.008 | 0.055 |
| $\rho$ | -0.126 | -0.017 | 0.107 |

Supplementary Table 4. Recovery analysis of the PF-HMM with response-level perseveration. The table shows summary statistics of errors between true (simulated) parameters and those recovered by the model. The 25%, 50% (Median), and 75% quantiles of the error distributions are reported. Both belief update parameters about stochasticity ($\sigma_s$) and volatility ($\sigma_v$) show median recovery errors close to zero, indicating effective recovery of parameters with no systematic bias.

| | $PLV$ | $PLM$ | $V$ | $S$ | $Acc$ | $I$ |
|---|---|---|---|---|---|---|
| Coeff. | -0.015 | 0.002 | -0.011 | 0.002 | 0.000 | 0.030 |
| S.E.M. | 0.004 | 0.002 | 0.009 | 0.006 | 0.001 | 0.002 |
| t-value | -3.533 | 0.945 | -1.239 | 0.423 | -0.579 | 15.009 |
| p-value | 0.001 | 0.348 | 0.219 | 0.673 | 0.564 | <0.001 |

Supplementary Table 5: Linear regression results for response times in the sea lion task using PF-HMM. Model-derived regressors were generated by first fitting the PF-HMM to participants' data and then using the median fitted parameters across subjects to produce trial-by-trial latent signals. Each column corresponds to a regressor included in the subject-level generalized linear model: $PLV$ (particle likelihood variability, quantified as the standard deviation of per-particle likelihoods), $PLM$ (mean per-particle likelihood), $V$ (estimated volatility), $S$ (estimated stochasticity), $Acc$ (binary indicator of whether the response matched the outcome), and $I$ (intercept). For each regressor, the table reports the group-level mean coefficient, standard error of the mean (S.E.M.), t-statistic, and p-value from a one-sample t-test across subjects. The significant negative coefficient for $PLV$ indicates that responses were slower when particle variability was low, consistent with increased difficulty in identifying the source of noise under particle indistinguishability.

| | $\delta$ | $\delta \times S$ | $\delta \times V$ | $\delta \times S \times V$ | $S$ | $V$ | $S \times V$ | $I$ |
|---|---|---|---|---|---|---|---|---|
| Coeff. | 1.014 | -0.110 | 0.216 | 0.021 | 0.002 | 0.005 | -0.010 | -0.010 |
| S.E.M. | 0.079 | 0.032 | 0.038 | 0.030 | 0.002 | 0.002 | 0.002 | 0.002 |
| t-value | 12.880 | -3.492 | 5.666 | 0.704 | 1.093 | 2.447 | -3.903 | -5.197 |
| p-value | <0.001 | 0.002 | <0.001 | 0.487 | 0.283 | 0.021 | 0.001 | <0.001 |

Supplementary Table 6. Fitted blockwise HMM to turtle task data (n = 30). This table presents the mixed-effects regression results for all regressors. The columns list each regressor with corresponding coefficient estimates (Coeff.), standard errors of the mean (S.E.M.), t-values, and p-values form two-sided t-tests. The term $\delta$ is defined as the observation minus previous belief ($\delta_t = o_t - r_{t-1}$). The factors $S$ and $V$ refer to the true stochasticity and true volatility, respectively, while $I$ represents the intercept term. Interaction terms (e.g., $\delta \times V, \delta \times S$, etc.) capture interaction effects of $\delta$ with the true environment parameters.

| | $PLV$ | $PLM$ | $V$ | $S$ | $Acc$ | $I$ |
|---|---|---|---|---|---|---|
| Coeff. | -0.032 | -0.006 | -0.009 | 0.014 | 0.005 | 0.046 |
| S.E.M. | 0.011 | 0.003 | 0.012 | 0.007 | 0.001 | 0.005 |
| t-value | -2.799 | -1.894 | -0.740 | 2.077 | 4.386 | 8.449 |
| p-value | 0.009 | 0.068 | 0.465 | 0.047 | <0.001 | <0.001 |

Supplementary Table 7: Linear regression results for response times in the turtle task using PF-HMM. Model-derived regressors were generated by first fitting the PF-HMM to participants' data and then using the median fitted parameters across subjects to produce trial-by-trial latent signals. Each column corresponds to a regressor included in the subject-level generalized linear model: $PLV$ (particle likelihood variability, quantified as the standard deviation of per-particle likelihoods), $PLM$ (mean per-particle likelihood), $V$ (estimated volatility), $S$ (estimated stochasticity), $Acc$ (binary indicator of whether the response matched the outcome), and $I$ (intercept). For each regressor, the table reports the group-level mean coefficient, standard error of the mean (S.E.M.), t-statistic, and p-value from a one-sample t-test across subjects. The significant negative coefficient for $PLV$ indicates that responses were slower when particle variability was low, consistent with increased difficulty in identifying the source of noise under particle indistinguishability.

| Models | MF | PXP |
| --- | --- | --- |
| PFHMM | 0.411 | 0.388 |
| PFHMM + $\rho$ | 0.589 | 0.612 |

Supplementary Table 8. Bayesian model comparison across PF-HMM and PF-HMM with response-level perseveration parameter ($\rho$). The table reports model frequency (MF) and protected exceedance probability (PXP) obtained from group-level Bayesian model selection. Including the perseveration parameter modestly improves model evidence, indicating that accounting for response-level biases enhances explanatory adequacy without altering the core inference structure of the model.

| | Sea lion task | | | Turtle task | | |
|---|---|---|---|---|---|---|
| | 25% quantile | Median | 75% quantile | 25% quantile | Median | 75% quantile |
| $\sigma_s$ | 0.009 | 0.092 | 0.420 | 0.014 | 0.100 | 0.352 |
| $\sigma_v$ | 0.001 | 0.042 | 0.202 | 0.001 | 0.060 | 0.253 |
| $\rho$ | 0.164 | 0.517 | 1.117 | 0.232 | 0.665 | 0.927 |

Supplementary Table 9. Summary statistics of PF-HMM-fitted parameters with response-level perseveration parameter. The three columns show the 25%, 50% and 75% quantiles for the fitted diffusion parameters controlling belief updates about stochasticity ($\sigma_s$) and volatility ($\sigma_v$), as well as the perseveration parameter ($\rho$). The perseveration parameter captures response-level bias to repeat or switch the previous choice and operates independently of latent state inference. Including this parameter improves model fit while leaving the qualitative dissociation between volatility- and stochasticity-related learning effects unchanged.

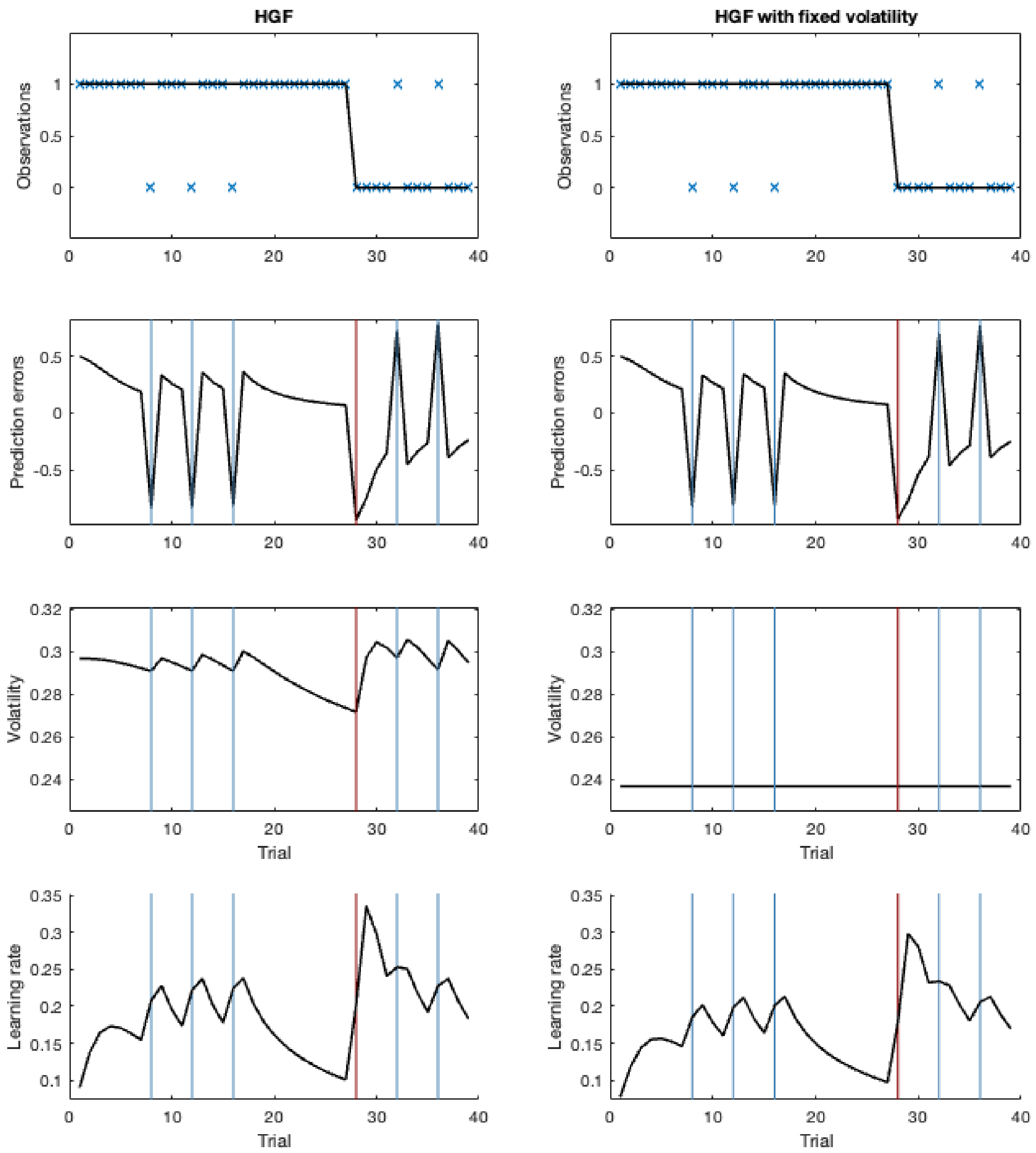


Supplementary Figure 1. Learning rate signal for HGF. Left column: Standard HGF fitted to the task. Right column: HGF with volatility fixed, preventing volatility-driven changes in the learning rate. From top to bottom: outcomes and hidden state, prediction errors, volatility, and learning rate. Across both conditions, learning rate increases sharply following large prediction errors, even when surprises are entirely due to stochasticity and volatility is held constant (right column). This illustrates that in sigmoid–Kalman models, the learning rate is intrinsically contaminated by the magnitude of prediction error, independent of volatility. Thus, apparent learning-rate increases does not reflect inferred volatility, but instead arise from nonlinear effects of the sigmoid transformation, as formally shown in Appendix B.

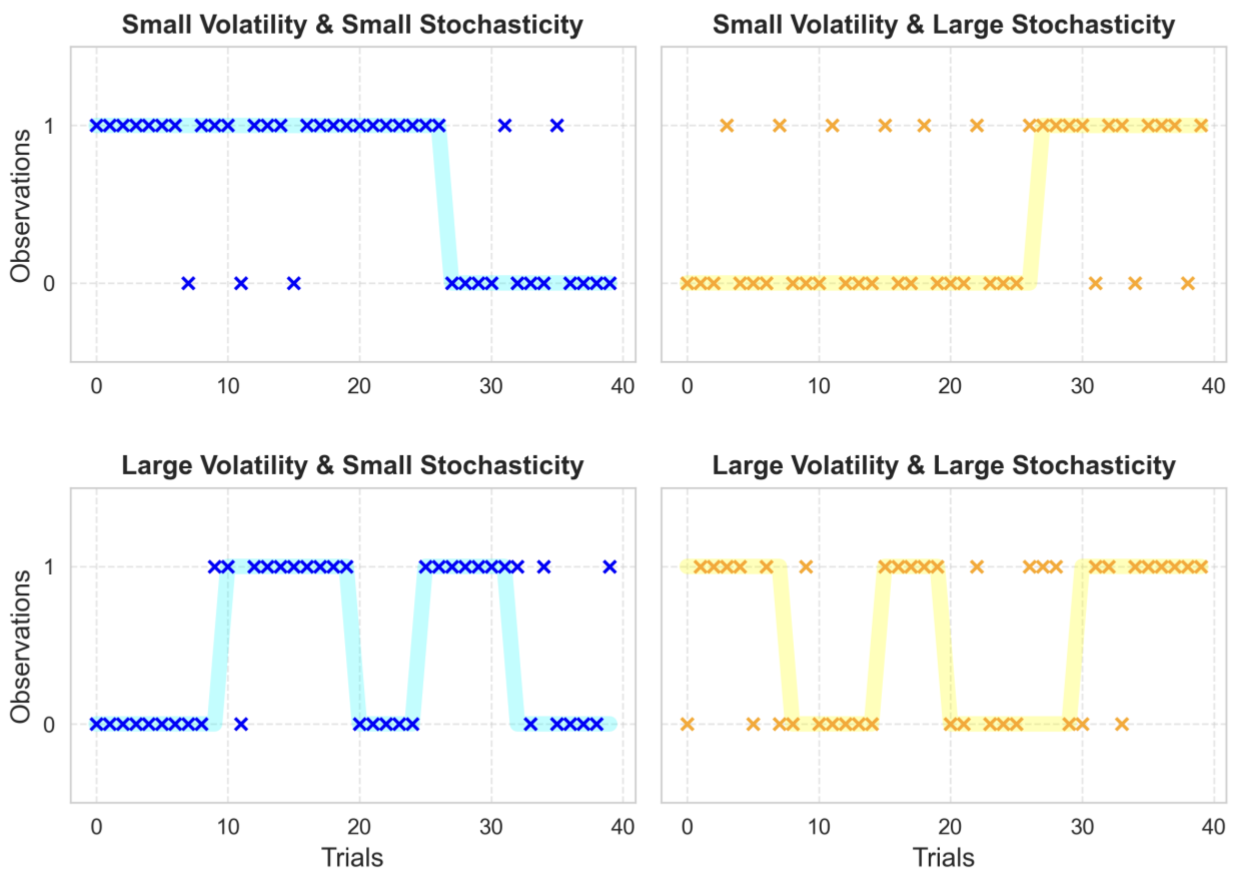


Supplementary Figure 2. Illustration of actual timeseries used in the sea lion task. The solid lines represent the hidden state, whose variability depends on volatility levels. Individual data points are marked separately, with their variations relative to the hidden state determined by the stochasticity level.

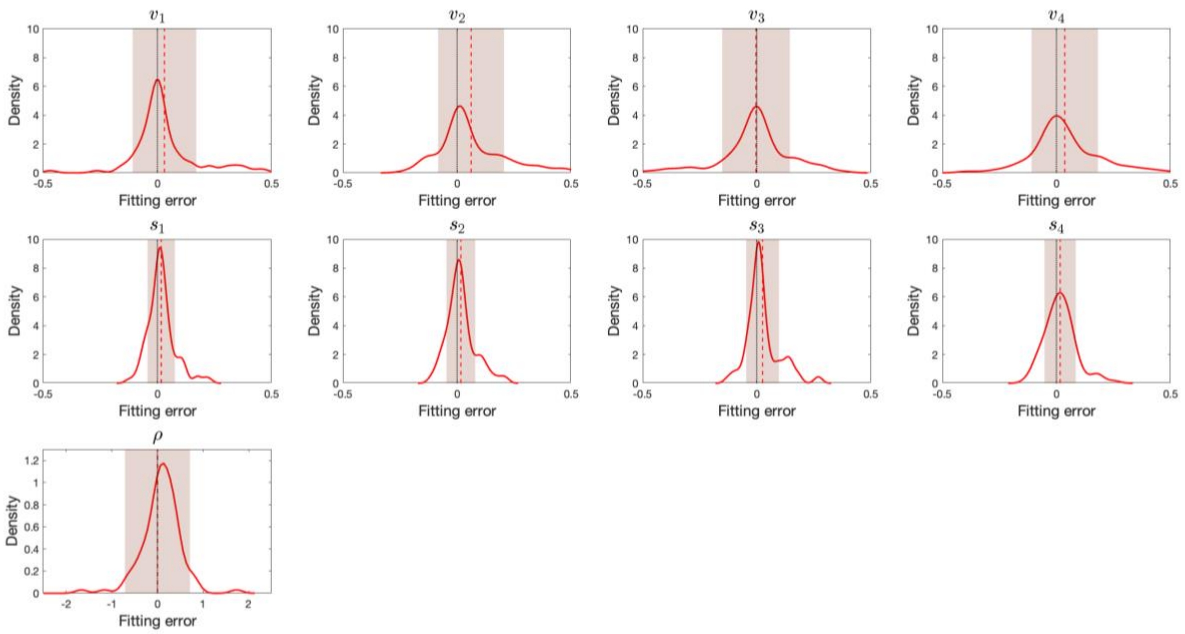


Supplementary Figure 3. Empirical distributions of fitting errors in the blockwise HMM with response-level perseveration recovery analysis. Each panel shows the kernel density estimate of the fitting error (red curve), defined as the true parameter minus the recovered parameter. The red vertical line indicates the mean fitting error, and the shaded region marks ±1 standard deviation around the mean. The black vertical line at zero denotes perfect recovery. Recovery was evaluated using 100 simulated datasets. For each simulation, raw HMM parameters (four volatility, four stochasticity, and one perseveration parameter) were sampled from a Gaussian distribution whose mean and standard deviation were estimated from the empirically fitted parameters of the real sealion task. The raw volatility and stochasticity parameters were transformed via a half-sigmoid mapping to constrain them to the interval (0, 0.5), while the perseveration parameter was left in raw space. Narrower distributions centered near zero indicate higher recovery accuracy. The results show that parameter estimates are unbiased, with mean fitting error close to zero. See also Supplementary Table 1.

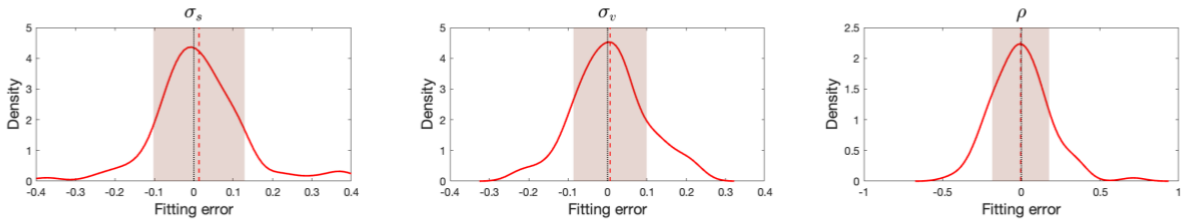


Supplementary Figure 4. Empirical distributions of fitting errors in the PF-HMM with response-level perseveration recovery analysis. Each panel shows the kernel density estimate of the fitting error (red curve), defined as the true parameter minus the recovered parameter. The red vertical line indicates the mean fitting error, and the shaded region represents ±1 standard deviation around the mean. The black vertical line at zero denotes perfect recovery. Recovery was evaluated using 100 simulated datasets. For each simulation, true parameters were sampled independently from uniform distributions: Narrow distributions centered near zero indicate accurate and unbiased parameter recovery. See also Supplementary Table 4.

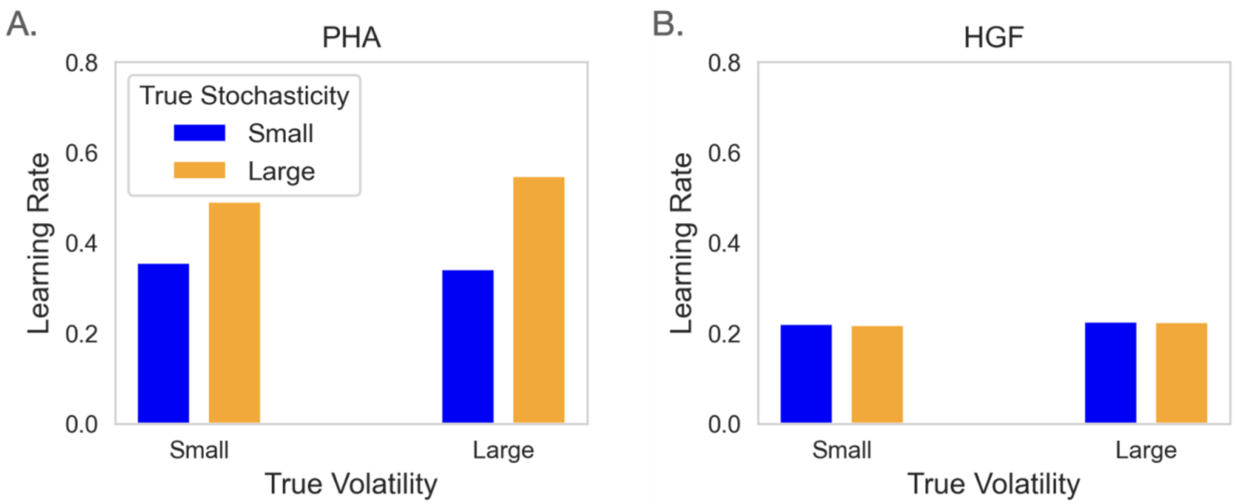


Supplementary Figure 5. Learning rates from alternative models using sea lion task data. (A) PHA, and (B) HGF. Parameters were fitted individually, and the median fitted parameters across participants were used to generate belief trajectories. Learning rate for both models were calculated using the same regression method presented in the main text. Unlike the PF-HMM, which predicts increased learning under high volatility and decreased learning under high stochasticity, both models fail to reproduce the opposing volatility–stochasticity pattern observed in the behavioral data.
